# Sustained Antigen Stimulation to Evoke and Study Negative feedback Systems responsible for Self-Tolerance/Tumor Immune Escape and transition to the M2 macrophage

**DOI:** 10.1101/2025.04.13.648563

**Authors:** Elizabeth A. Mazzio, Andrew S Barnes, Ramesh B Badisa, Selina F Darling-Reed, Karam FA Soliman

## Abstract

Chronic inflammation plays an obscure role in cancer initiation, with broad references implicating immune exhaustion (IEX) or free radical mediated cell damage. Chronic inflammation is, however, paradoxically synonymous with the term “immune tolerance” which in other cases presents itself as a therapeutic limitation to the efficacy of tumor immune therapies particularly those involving microbial -associated molecular patterns (MAMPs) in regimens. As “tolerance” remains to this day a “phenomenon” there is a pressing need to fully understand every biological aspect of this apparent negative feedback response, because doing so will serve to guide development of targeted immune therapies. In this work, we employ a rudimentary model, which can be adopted in which it is possible to provoke negative feedback through sustained antigen stimulation in immunocompetent cells, as we monitor, define and characterize phenotype evolution using next generation whole transcriptome sequencing + validation studies. This model can be used to study /create *in vitro* the “M2” phenotype, which is itself involved in aggressive tumors with synergistic rapid expansion of myeloid-derived suppressor cells (MDSCs), dysfunctional CD8+ T cytotoxic (CTL)/ and dysfunctional natural killer (NK) cells. Briefly, the data in this work, shows negative feedback dominates at 7 to 11 days, after acute being associated with phase specific (time dependent) elevated checkpoints; e.g. PDL1+/MSN, HAVCR2/TIM-3+, SPP1+, C3ar1+, CD73+, IL1RN+, LILRbs+, glycoproteins, integrins etc. Here we report on phase specific patterns and bidirectional changes aligning with immune escape, much centered around loss of host defense against viral infection and malignancy. Negative feedback is associated with rampant induction of degradative proteases, SOCS/JAK/STAT/IL-10, the CCL2/7 axis tantamount to sustained loss of MHC1/2 antigen recognition systems, Type I interferon response, NOD signaling, antiviral/antibacterial defense, p62/SQSTM also aligning with a disturbed metabolic signature. The data in this work demonstrates that the colloquial terms “tolerance and IEX” are somewhat flawed terminology, because this negative feedback is a potent, intentional and formidable immune offense to eradicate the active arm of immune defense.

**Abstract Truncated /Removed due to word count:** The data show that “tolerance” aligns with six distinct chronological differential gene (DEG) expression patterns that circumscribe extensive immune suppression which dominate during chronic inflammation. These involve the following time dependent patterns: 1) transcripts overexpressed, e.g. checkpoint receptors; PDL1+/MSN, HAVCR2/TIM-3+, SPP1+, C3ar1+, CD73+, IL1RN+, LILRbs+, glycoproteins, etc.; 2) transcripts overexpressed only in chronic; e.g cytokine suppressor signaling (SOC3/Jak/Stat), cyto/chemokines (IL-10, CCL2, CCL7), proteases (cathepsins L, D, K, Adam 8, PIM2), and adhesives (TSPAN3, QSOX1, PDPN, ITGA5), (PLK2, ADGRE1, CALM1, PCNA, etc.); 3) transcripts downregulated in acute and chronic; e.g. a severe collective loss in MHC1/II antigen presentation capacity (CD74, H2-Q4, H2-Q6, etc.), NOD signaling, and interferon (IFN) type I signaling systems; 4) transcripts downregulated in chronic only, including OXPHOS/metabolic genes (Aldo A, C, Eno2, Gpi1, etc.), antiviral/antibacterial defense genes (Lyz1, Lyz2, Card19, Ninj1), and autophagy-related genes (p62/SQSTM1). In the category of Tolerance were: 5) transcripts induced sharply in acute, no longer responsive in chronic; IFN Type 1 antiviral response genes (OAS, BST2, ISG15, ISG20, IRF7, RSAD2/Viperin), TLR2, antibacterial defense genes (SAA3, SP140), and proinflammatory cytokines (CCL5, TNF, IL1a, and IL1b) along with the IL-1/TLR signaling axis. Last, 6) reverse tolerance, corresponded to restored baseline levels to maintain cell mitotic and thymosin homeostasis. In conclusion, these data suggest chronic inflammation precipitates negative feedback, aligning with the same checkpoint targets being sought after today in tumor immune therapies.

## Introduction

For over 100 yrs, there continues to remain an obscure, cryptic connection involving host response to bacterial infection and malignant self-cells. First, long forgotten were the earlier accounts by physicians Busch, Coley, and Fehleisen reporting curative spontaneous remission of human cancers being observed in individuals post-acute febrile infection (38-40°C) infected with Streptococcus pyogenes. (1) While therapeutic administration of inactivated bacteria (Coleys Toxin) was the first tumor immune therapy, it had moderate success. The active immune boosting antigen later identified as gram-negative cell wall LPS, was then characterized as a microbial-associated molecular pattern (MAMP). Today MAMPs are used in vaccines and tumor immune therapies e.g. Bacillus Calmette-Guerin (BCG) for bladder cancer, while synthetic LPS analogues tested in clinical trials – have met with moderate success, with limited therapeutic efficacy arising from “tolerance” (2–5) also known as “immune exhaustion” (6). Again, there is great uncertainty as to the provocative role in chronic inflammation and cancer, and from the data in this work, we provide evidence that tolerance induced by chronic inflammation are one in the same. They both involve provocation of a formidable immune negative feedback response. Because tolerance to this day remains poorly understood (7); see Review (8), it is imperative that we unveil its mechanism because it is implicated in limited response to anticancer/adoptive immunotherapies (9–13) in addition to post septic immunosuppression,and subsequent rise to opportunistic infections which claim millions of lives annually (14, 15).

Further needed are creative models that clarify the double-edged sword that exists for “acute” vs “chronic” exposure to immune active bacterial antigens. On the one hand acute (high dose short term) MAMP activation of toll-like receptors (TLRs), can provoke a cytokine storm, sepsis, multiorgan failure, and death and on the other hand the same is commonly used in vaccines and tumor immune therapies (16). That said, there is vast distinction between acute (immune defense/cytokine storm) and chronic (negative feedback). Chronic inflammation and its involvement in cancer while still obscure, becomes more complicated as we learn about the power of the tumor microbiome/commensal bacteria (17) which can originate from inflammatory pathogenic microbes (e.g. the oral cavity such as *Porphyromonas gingivalis),* with ability to travel systemically and end up in a “cancer microbiome” inextricably linked to immune escape in the tumor microenvironment (TME). Therefore, the ultimate question seems to be at this point, does cancer use chronic inflammation as a mechanism to reinforce negative feedback creating the very tumor immune escape we seek to fight against? Much of the evidence in the literature and in the data in this work, suggests this concept could be highly plausible.

Even further, there is a need for creative new models which can re-create the loss of immune surveillance *in vitro*. The original premise of this work was only to elucidate the intricacies of tolerance, and define IEX, however the findings, unearthed a roadmap to hundreds of checkpoints, bidirectional and time dependent. In this model, while rudimentary in sophistication, seeks to investigate phenotype evolution from resting, to acute and through to chronic by long term sustained inflammatory activation by LPS/TLR4/MyD88 signaling, in macrophages, it is noteworthy that this type of study requires rigorous controls to prevent, artifact / false variable influence by the biological products induced from sustained antigen activation. The outcome of this model abides in the known time line of refractory immune suppression that occurs post sepsis; starting with acute phase (cytokine storm/ LPS/ septic shock) 1-3 days, followed by severe immunosuppression 7-11 days (negative feedback), the later involving the same reported transformation of macrophages from an M1 to an M2 subpopulation, as described in aggressive cancers (18).

The overall findings, provide 1) an adoptable model to recreate tumor immune suppression in the lab through sustained antigen stimulation of immunocompetent cells 2) a new understanding into the phenomenon of “tolerance” and “immune exhaustion” neither of which are accurate 3) and results that show an aggressive sustained negative feedback response is evoked through 8 bidirectional time specfic patterns. Moreover, these findings clearly show that while free radicals may be involved with initiation of cancer, chronic inflammation could evoke the very same immune escape by which cancer can take hold and thrive.

## Methods

### Cell culture

Mouse macrophage (RAW 264.7) RRID:CVCL0493 cells were purchased from American Type Culture Collection (Manassas, VA, USA). The cells were maintained in high-glucose DMEM [glucose 4500 mg/L] containing 5% FBS, 4 mM L-glutamine, and penicillin/streptomycin (100 U/0.1 mg/mL). The culture conditions were maintained at 37°C in a 5% CO_2_/atmosphere in preparation for subculture for experimentation. For the experiments, the medium consisted of DMEM (minus phenol red) [glucose 4500 mg/L], 4 mM L-glutamine, 5% FBS and penicillin/streptomycin (100 U/0.1 mg/mL). LPS from *E. coli* (O111:B4) was used in all the experiments throughout this work at 500 ng/mL (working).

### Endotoxin tolerance induction method validation

A pre-method validation study was conducted to ensure the lack of accumulated cellular byproducts in the media, particularly for LPS treatment. This involved determining the dose‒response effect on the cell density [50–50,000 cells/well /200 µL per well] in a 96-well plate treated with 500 ng/mL LPS for 24 h. Wells that produced <L.O.D. for NO at 24 h were chosen for extrapolated flask experiments (75 cm3), and intermittent media + LPS exchange was continued every 18 h, with the monitored absence of accumulated nitric oxide (which did not acquire tolerance, prevalidated). The cells at all time points were fully viable, collected, and labeled as follows: 1) negative control: control resting; 2) positive control: LPS-treated acutely at 500 ng/mL (24 h); 3) chronic 1; 7 d sustained 500 ng/mL (chronic) LPS; and 4) chronic 2; 11 d sustained 500 ng/mL (chronic) LPS. At the end of the study, the cell pellets and supernatants were collected and stored at −80°C.

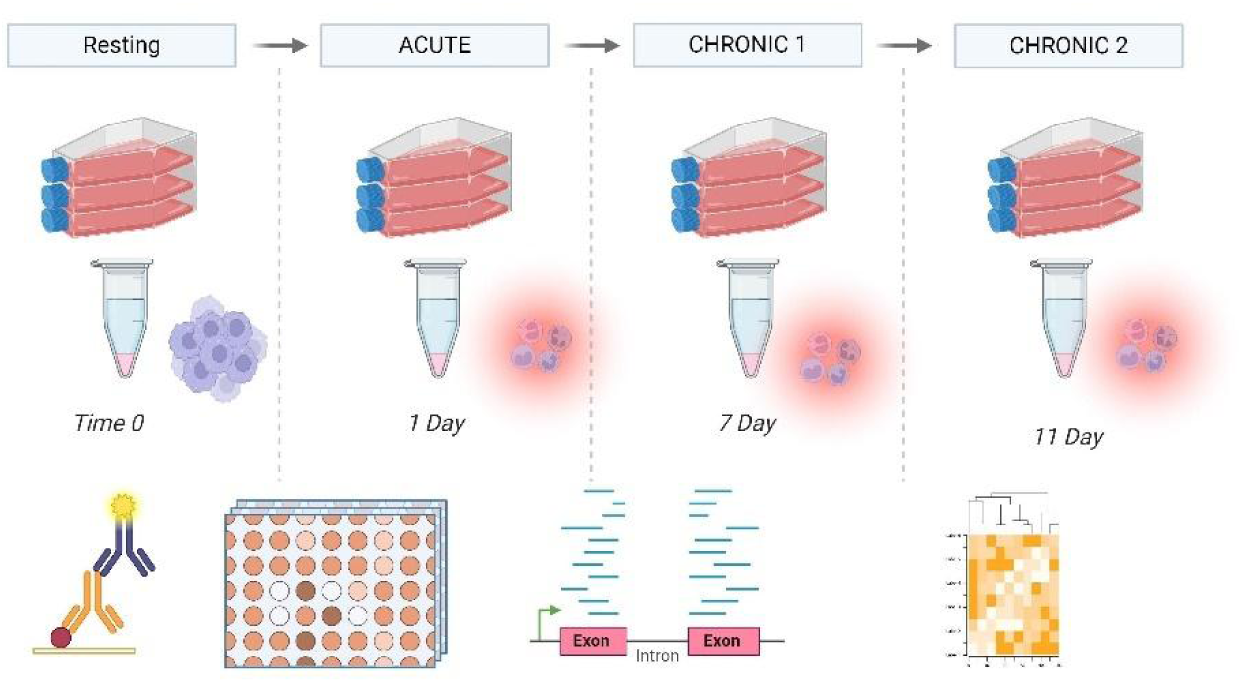

### Nitric oxide monitoring

The quantification of nitrite (NO2−) was performed via the use of the Greiss reagent. The Greiss reagent was prepared by mixing an equal volume of 1.0% sulfanilamide in 10% phosphoric acid and 0.1% N-(1-naphthyl)-ethylenediamine in deionized water, which was added directly to a sample of supernatant (experimental medium consisting of DMEM - phenol red) and incubated at room temperature for 10 min. Samples were analyzed at 540 nm via a Synergy HTX multimode reader (Bio-Tek; Winooski, VT, USA).

### RNA Sequencing

RNA sequencing was performed with the assistance of LC Biosciences (Houston, Texas, USA). Total RNA was extracted via TRIzol reagent (Thermo Fisher, CA, USA; cat. 15596018) following the manufacturer’s instructions. The total RNA quantity and purity were analyzed via a Bioanalyzer 2100 and RNA 6000 Nano LabChip Kit (Agilent, CA, USA; cat. 5067--1511); high-quality RNA samples with a RIN >7.0 were used to construct the sequencing library. After total RNA was extracted, it was purified from total RNA (5 µg) via Dynabeads Oligo (dT) (Thermo Fisher) via two rounds of purification. Following purification, the mRNA was fragmented into short fragments via divalent cations under elevated temperature [Magnesium RNA Fragmentation Module (NEB, USA, cat. e6150) at 94°C for 5–7 min]. The cleaved RNA fragments were then reverse transcribed to create cDNA by SuperScript™ II Reverse Transcriptase (Invitrogen, USA, cat. 1896649), which was subsequently used to synthesize U-labeled second-strand DNAs with *E. coli* DNA polymerase I (NEB, cat. m0209), RNase H (NEB, cat. m0297) and dUTP Solution (Thermo Fisher, cat. R0133). An A-base was then added to the blunt ends of each strand, preparing them for ligation to the indexed adapters. Each adapter contained a T-base overhang for ligating the adapter to the A-tailed fragmented DNA. Dual-index adapters were ligated to the fragments, and size selection was performed with AMPureXP beads. After heat-labile UDG enzyme (NEB, cat. m0280) treatment of the U-labeled second-strand DNAs, the ligated products were amplified via PCR under the following conditions: initial denaturation at 95°C for 3 min; 8 cycles of denaturation at 98°C for 15 sec, annealing at 60°C for 15 sec, and extension at 72°C for 30 sec; and a final extension at 72°C for 5 min. The average insert size for the final cDNA libraries was 300±50 bp. Finally, we performed 2×150 bp paired-end sequencing (PE150) on an Illumina NovaSeq™ 6000 following the vendor’s recommended protocol.

### Bioinformatics analysis

Sequence and filtering of Clean Reads: A cDNA library was constructed from pooled RNA from murine species via technology and then sequenced with the Illumina 6000 platform. Using the Illumina paired-end RNA-seq approach, we sequenced the transcriptome, generating a total of one million 2 × 150 bp paired-end reads. The readings obtained included raw reads containing adapters or low-quality bases which affected the subsequent assembly and analysis. Thus, to obtain high quality clean reads, the reads were further filtered by Cutadapt RRID: SCR_011841 (https://cutadapt.readthedocs.io/en/stable/,version:cutadapt-1.9). The parameters were as follows: 1) reads containing adapters were removed; 2) reads containing polyA or polyG were removed; 3) reads containing more than 5% unknown nucleotides (N) were removed; and 4) low-quality reads containing more than 20% low quality (Q value≤20) bases were removed. The sequence quality was subsequently verified via FastQC (RRID:SCR_014583) (http://www.bioinformatics.babraham.ac.uk/projects/fastqc/,0. 11.9), including the Q20, Q30 and GC contents of the clean data. After that, a total of G bp of cleaned, paired-end reads were produced(19, 20). Alignment with the reference genome: We aligned the reads of all the samples to the mouse reference genome via the HISAT2 (RRID:SCR_015530) (https://daehwankimlab.github.io/hisat2/,version:hisat2-2.0.4) package, which initially removes a portion of the reads based on quality information accompanying each read and then maps the reads to the reference genome. HISAT2 allows multiple alignments per read (up to 20 by default) and a maximum of two mismatches when mapping the reads to the reference. HISAT2 builds a database of potential splice junctions and confirms these results by comparing the previously unmapped reads against the database of putative junctions(21, 22).

### Quantification of gene abundance

The mapped reads of each sample were assembled via StringTie (RRID:SCR_016323) (http://ccb.jhu.edu/software/stringtie/,version:stringtie-1.3.4d) with the default parameters. All the transcriptomes from all the samples were merged to reconstruct a comprehensive transcriptome via GffCompare software (http://ccb.jhu.edu/software/stringtie/gffcompare.shtml,version:gffcompare-0.9.8). After the final transcriptome was generated, StringTie and Ballgown (http://www.bioconductor.org/packages/release/bioc/html/ballgown.html) were used to estimate the expression levels of all the transcripts, and the expression abundance of the mRNAs was determined by calculating fragment per kilobase of transcript per million mapped reads (FPKM) value. Gene differential expression analysis was performed via DESeq2 software between two separate groups (and via edgeR between two samples). Genes whose false discovery rate (FDR) was less than 0.05 and absolute fold change ≥2 were considered differentially expressed genes (22–24). Alternative splicing (AS) analysis: rMATS (version 4.1.1) (http://rnaseq-mats.sourceforge.net) was used to identify alternative splicing events. We identified AS events with a false discovery rate (FDR) of p<0.05 in comparison with significant AS events. The classification of alternative splicing is as follows: SE: skipped exon MXE: mutually exclusive exon A5SS: alternative 5’ splice site A3SS: alternative 3’ splice site RI: retained intron (25, 26). All samples were run in triplicate. The data are available from the National Institutes of Health (NIH) Gene Expression Omnibus (RRID:SCR_005012); series number GSE263901.

### Protein Semiquantitative: Reference Check Mouse cytokine antibody array

Mouse cytokine antibody arrays (Ray Biotech, GA, USA; product code: AAM-CYT-1000) were used to profile the cytokine concentration in the supernatant for each sample set. Each experiment was performed in duplicate according to the manufacturer’s instructions. Briefly, antibody-coated array membranes were first incubated for 30 min with 1 mL of blocking buffer and then replaced with 1 mL of sample supernatant, after which the membranes were incubated overnight at 4°C with mild shaking. The medium was then removed, and the membranes were washed three times, and subsequently incubated with 1 mL of biotin-conjugated antibodies for 6 h. After repeated washes, the biotin-conjugated antibodies were removed. The membranes were incubated with HRP-conjugated streptavidin (2 h), and then evaluated via densitometry via a chemiluminescence substrate on a Versa Doc Imager/Quantity One software from Bio-Rad (Hercules, CA, USA).

### Protein reference check (ELISA)

CCL5 was entirely subject to tolerance, whereas chronic inflammation and resting conditions were similar. Compared with acute elevation, chronic LPS treatment resulted in a [-8.83 Log2FC p-Val=3.00E-303] reduction. CCL5 protein was validated via the RayBio® Mouse RANTES (CCL5) ELISA. Cell supernatants from all groups were evaluated, and assays were performed in accordance with the manufacturer’s guidelines. The samples were analyzed at 450 nm via a BioTek Synergy HTX Multimode Reader (Agilent Technologies; Santa Clara, CA, USA).

### Data analysis

The bioinformatics tools used for the analysis of the data in this work required the use of various platforms, including the Kanehisa Laboratories (KEGG) (RRID: SCR_012773) (computational pathways (27, 28), Reactome gene sets(29), DAVID (RRID: SCR_001881) functional ontology (30, 31) and Enricher KG (32, 33). Basic statistical analysis was performed via Graph Pad Prism (RRID: SCR_002798) (version 3.0; Graph Pad Software Inc. San Diego, CA, USA), with the significance of differences between the groups assessed via one-way ANOVA, followed by Tukey’s post hoc means comparison test, or Student’s t test.

## Results

The first and most important analysis classified all the changes according to their distinct DEG dynamic patterns. These patterns were self-evident by RNA-seq data from resting to acute (24 hours) and acute to chronic inflammation (7 and 11 days), and many failed to meet the true definition of tolerance. These patterns are presented in a representative graph (**Figure 1)**. Figure 1 (Top panel) displays patterns that lead to chronic sustained “overexpressed” genes at days 7 and 11, including those transcripts that are overexpressed during acute inflammation and sustained in chronic inflammation (acute/sustained in chronic inflammation), transcripts that are overexpressed in chronic inflammation only, and transcripts that are slightly attenuated from 24-hour acute LPS but remain above the resting state during chronic inflammation. The last category involves reverse tolerance whereby acute LPS involves repression, an effect that is lost in chronic inflammation and returns to the resting baseline. **Figure** 1 (lower panel) shows patterns leading to sustained repression of transcripts in chronic inflammation. These include transcripts repressed in both acute and chronic conditions (days 7 and 11), repressed in chronic conditions only, transcripts subject to tolerance (elevated in acute conditions and lost in chronic conditions), repressed in acute conditions, and only mildly elevated in chronic conditions but remaining well below the resting cell control baseline. All future data will now be presented in terms of directional shifts such as “overexpressed” or “underexpressed” in chronic inflammation relative to the appropriate control group, which are both extremely important, while most research is focused on the few overexpressed checkpoints for immune therapy.

**Figure 1.**
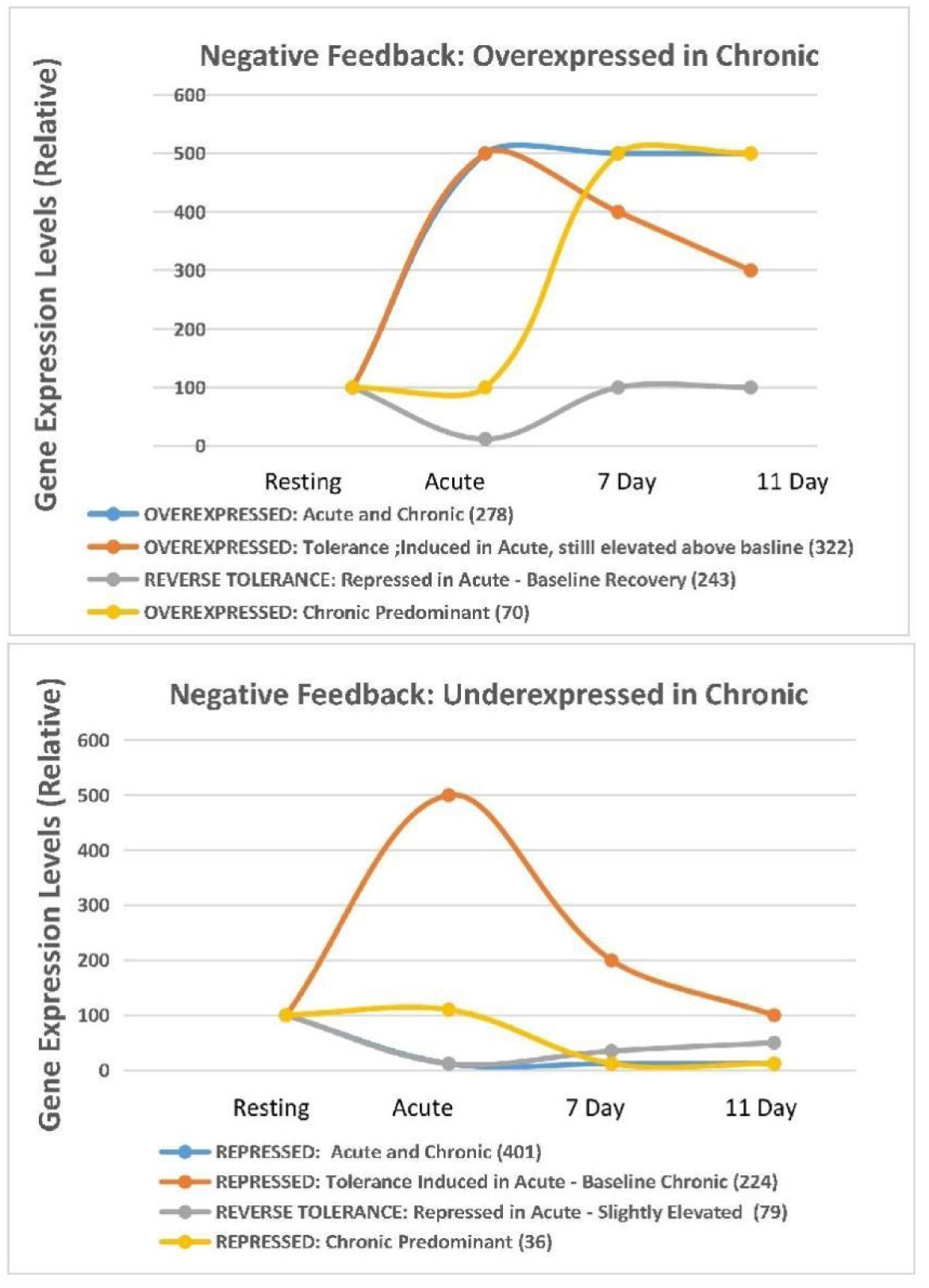
Chronological Transcript Changes associated with Chronic Inflammation; Resting, Acute and Chronic. The data shows pattern differential expressed genes (DEG) by # that fall in into 2 categories, and 8 subcategories. DEGS eventually over expressed (Top Panel) or under expressed (Bottom Panel) by 7-11 days. Classifications in the Top Panel shows transcripts a) up regulated in acute and sustained in chronic, b) up regulated in acute only c) up regulated in acute then subject to diminished response “tolerance” but remaining significantly higher than resting, and d) subject to full tolerance (up regulated in acute-no longer responsive). Likewise, the lower panel shows DEGs a) downregulated in acute and suppressed in chronic b), downregulated in chronic only, c) downregulated in acute with partial restoration “ tolerance” still not reaching resting baseline and d) subject to tolerance (downregulated in acute), returning to baseline in chronic. Significant transcript changes by number are reflected as (#) in the graphs.

With limitations in current bioinformatics methods, which are known, significant prominent changes in individual genes for which little is known cannot be detected. However, heatmaps generated from the input data of the chronically overexpressed DEGs (7,11 days) are presented in **Figure 2A** whereas those generated from the underexpressed DEGs are presented in Figure 2B via the Reactome ontology bioinformatics platform. These data show that chronic inflammation is associated with the overexpression of IL-10, the activation of the CD163 anti-inflammatory response, the SOCS Jak/Stat pathway, and CSF3 signaling with the repression of type I IFN signaling and proinflammatory inflammasomes. This heatmap is broad; therefore, we listed values for Log2FC and significance for DEGs, as presented in the legend of Figure 2. With the use of a broad scatterplot of Reactome gene sets **(Figure 3A),** all sequencing data were pooled into one plot according to functional clusters. Figure 3 -The top panel represents under expressed DEGs dominated by larger dominant clusters, whereas Figure 3-the bottom panel represents overexpressed DEGs dominated by larger dominant clusters. While observational comparative analysis is visually apparent, we provide a table **(Figure 3B)** to interpret the bidirectional events associated with chronic inflammation easily. In brief, the data from this analysis show that chronic inflammation triggers a collective loss in the host cytosolic sensor capacity of pathogen ssDNA, rRNA processing, and TICAM1-activation of IF3/IRF7 signaling. Moreover, FCGR3A-IL10 synthesis, reactive oxygen species (ROS), reactive nitrogen species (RNS), and selective autophagy lead to a switch toward a post replicative state, alterations in carbohydrate metabolism, protein degradation, insulin receptor signaling and lipoprotein remodeling. These broad changes will be clarified and integrated with greater specificity throughout the results and elaborated upon in the discussion.

**Figure 2A.**
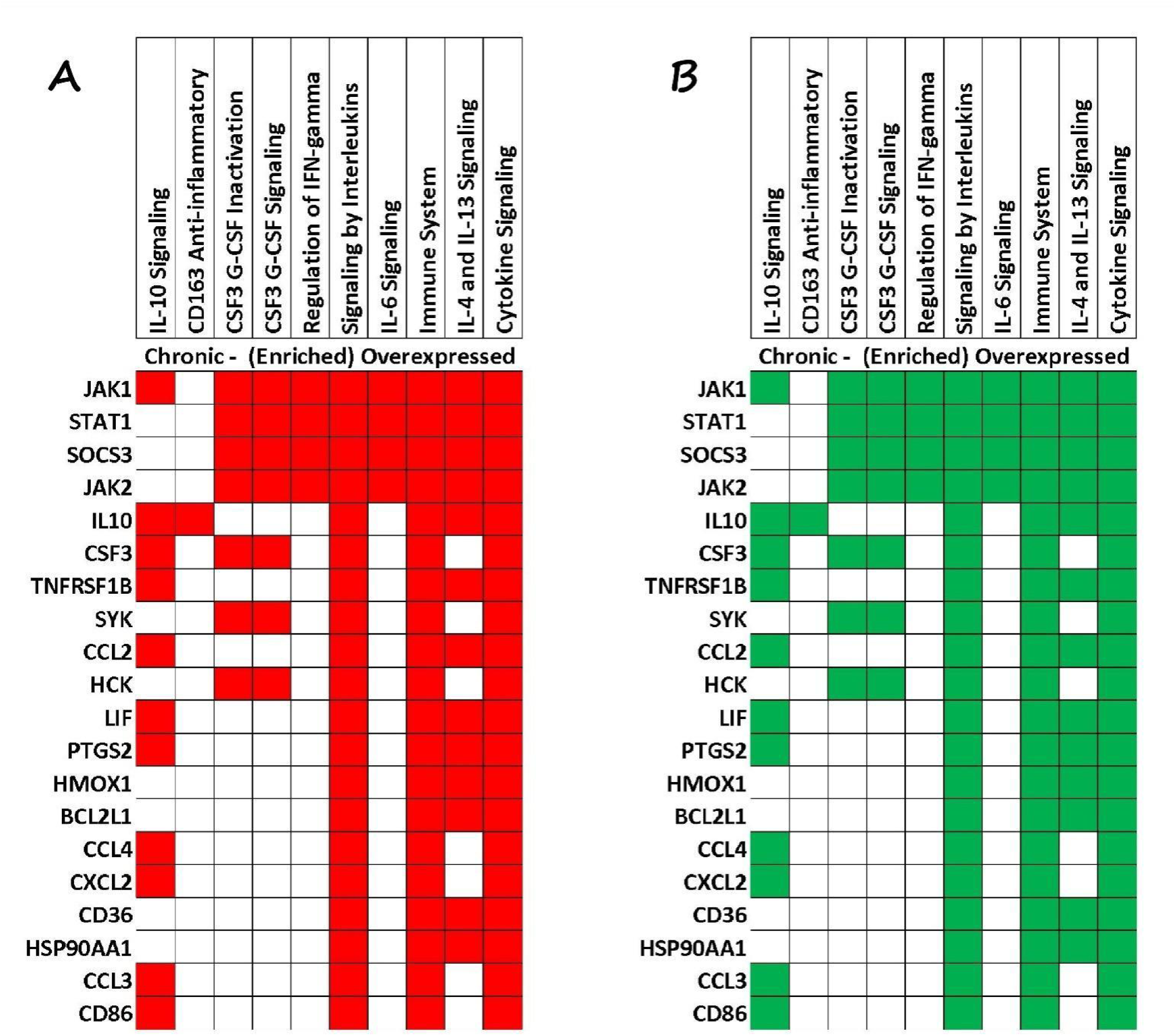
Down-regulated transcripts associated with chronic inflammation at 7-11 Days (Red/ Left Panel). Basic functional annotation of gene sets associated with down regulated processes during chronic inflammation represented by genes (X-axis) and pathways (Y-Axis) derived from Reactome bio-informatics. Genes under expressed correlate to losses in Type I IFN signaling, host defense against viral and bacterial pathogens, IL1α, IL1β, the inflammasome and antigen presentation. Major losses by Gene,FC and p-Value were observed for (\***Nfkb2** −2.01 Log2FC p-Val=6.37E-89) (**\*Isg15** −5.56 Log2FC p-Val<1.00E-300), (**\*Isg20** −3.71 Log2FC p-Val =1.91E-31) (**\*Il1a** −5.16 Log2FC p-Val <1.00E-300), (**\*Il1b** −4.49 Log2FCp-Val<1.00E-300), (**\*Gbp2** −2.99 Log2FC p-Val<1.00E-300), (**\*Bst2** −2.00 Log2FC p-VaL=3.86E-32), (**\*Ifit1** −6.31 Log2FCp-Val<1.00E-300), (**\*Ifit3** −4.97 Log2FC p-Val<1.00E-300), (**\*Ifit2** −2.64 Log2FC p-Val=3.09E-101), (**\*Oas3** −2.51 Log2FC p-Val<1=0.00E-300), (**\*Oas1g** −3.39 Log2FC p-Val=9.02E-32), (**\*Irf7** −4.82 Log2FC p-Val =4.24E-72), (**\*Rsad2** −4.13 Log2FC p-Val<1.00E-300); Additional prominent DEGs of note not on heatmap,(**Cd74** −4.2 Log2FC p-Val 5.72E-10),(**Fxyd2** −8.5 Log2FC p-Val =4.02E-12), (**Lsp1** −4.5 Log2FC p-Val=7.07E-11), (**\*Ifi27l2a** −3.4 Log2FC p-Val 1.79E-05), (**\*H2-Q6** −2.15 Log2FC p-Val=3.96E-218), (**\*H2-T24** −2.60 Log2FC p-Val=1.66E-73), (**\*H2-T23** −2.37 Log2FC p-Val=4.18E-69),(**\*H2-M2** −4.87 Log2FCp-Val=2.72E-53),(**\*H2-T23** −3.10 Log2FC p-Val=6.45E-47),(**\*H2-T10** −2.78 Log2FC p-Val=2.22E-14),(**\*H2-K2** −2.01 Log2FC p-Val = 1) 86E-12),(**\*H2-K1** −17.02 Log2FCp-Val=5.57E-10). **Figure 2B. Up-regulated transcripts associated with chronic inflammation at 7-11 Days (Green/ Right Panel).**Functional annotation of gene sets by heat map depicting Gene (X-axis) and Pathways (Y-Axis) analyzed using Reactome bio-informatics. DEGs associated with chronic inflammation involve up regulation of SOC3 Jak/Stat signaling, the CD163 anti-inflammatory response and CCL chemokines; These include by gene, FC, and p-Values (e.g.**Socs3** +4.9 Log2FC p-Val 8.73E-20), (**IL-10** +7.6 Log2FC; p-Val =2.8E-14), (**CCL2** +6.2 Log2FC; p-Val=8.5E-12),(**Ccl4** +2.1 Log2FCp-Val=1.63E-03), (**Lif** +3.7 Log2FC p-Val 3.57E-08); additional prominent DEGs of note not on heatmap; (**Klk9** +7.2 Log2FC; p-Val =7.0E-10),(**NOS2** +5.5 Log2FC; p-Val =5.7E-34),(**CCL7** +6.0 Log2FC; p-Val =5.4xE-11),(**Il1f9** +7.5 Log2FC; p-Val =1.14xE-27),(**S100A8** +7.3 Log2FC p-Val =4.3E-15) (**Csta3** +6.9 Log2FC p-Val =8.53E-08) (**Il33** +7.1 Log2FC p-Val =9.10E-040), (**Ccl12** +5.8 Log2FC p-Val =1.36E-03), (**Edn1** +5.7 Log2FC p-Val =6.74E-06), (**Il1f9** +7.5 Log2FC p-Val=1.14E-27), (**Sema3e** +5.7 Log2FC p-Val =9.59E-19), (**Urah** +6.7 Log2FC p-Val =6.22E-04), (**Vav1** +18.7 Log2FC p-Val=1.74E-19), (**Mast3** +11.5 Log2FC p-Val 6.12E-10).

**Figure 3A.**
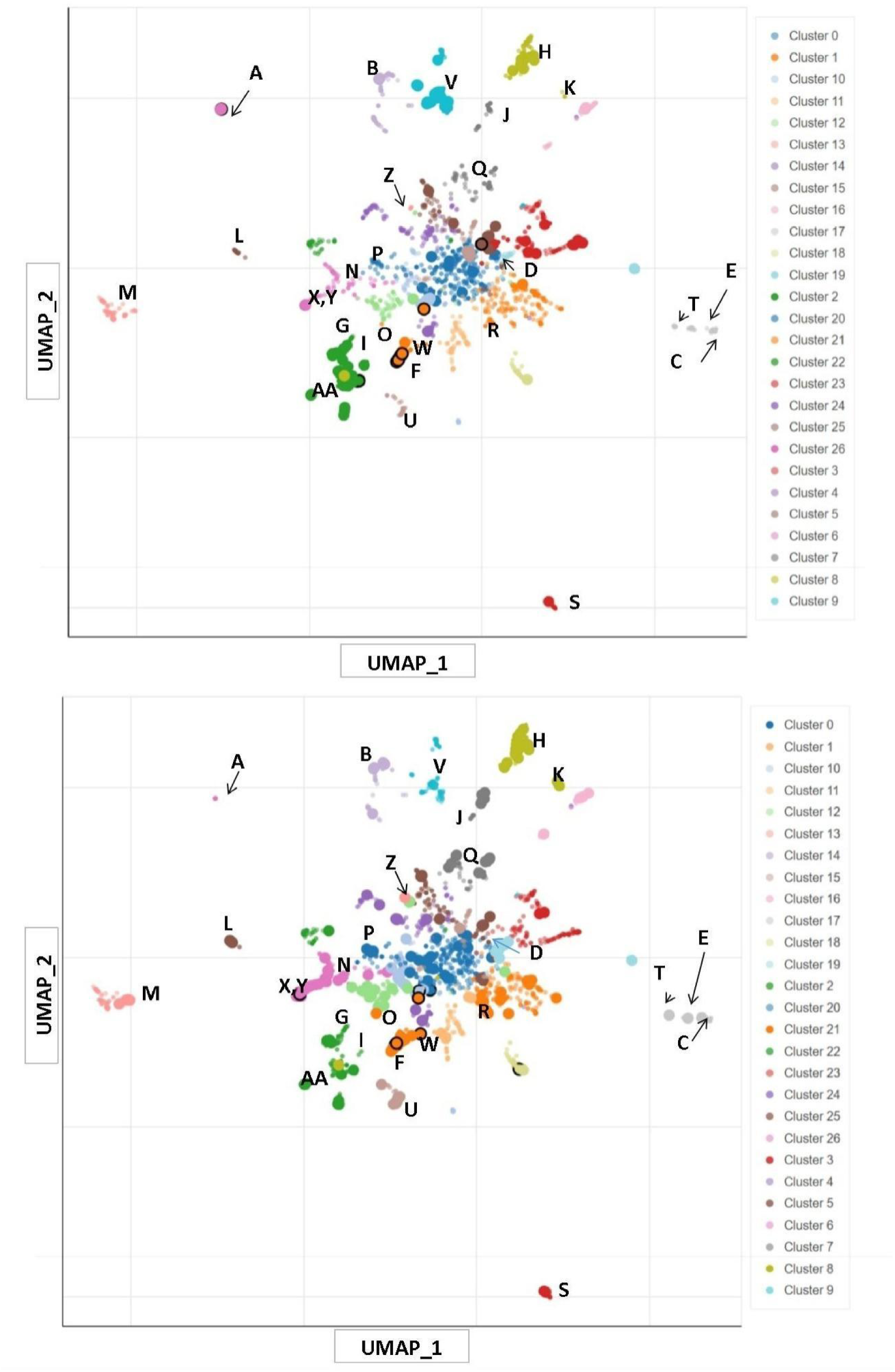
Bi-directional Functional Clustering of cell systems impacted by chronic inflammation at 7-11 Days. Functional clusters altered in chronic inflammation are presented for both both the up and down directions and correlate to alphabetized tabulations in Figure 3B. Each term represents a system defined in the Reactome library where term frequency-inverse document frequency (TF-IDF) values were computed for the Reactome gene set and a UMAP was applied to the resulting values. Generally, terms with more similar gene sets are positioned closer together. The terms are colored according to the automatically identified clusters computed with the Leiden algorithm applied to the TF-IDF values. The darker and larger the point is, the more significantly enriched the term for the described condition. The scatter plot of the top panel (represents under expressed in chronic) and the bottom panel (over expressed in chronic), with the larger spots dominating changes. For ease and clarity of interpretation– we have provided the following Figure (Tabulated) 3B for reference.

**Figure 3B.**
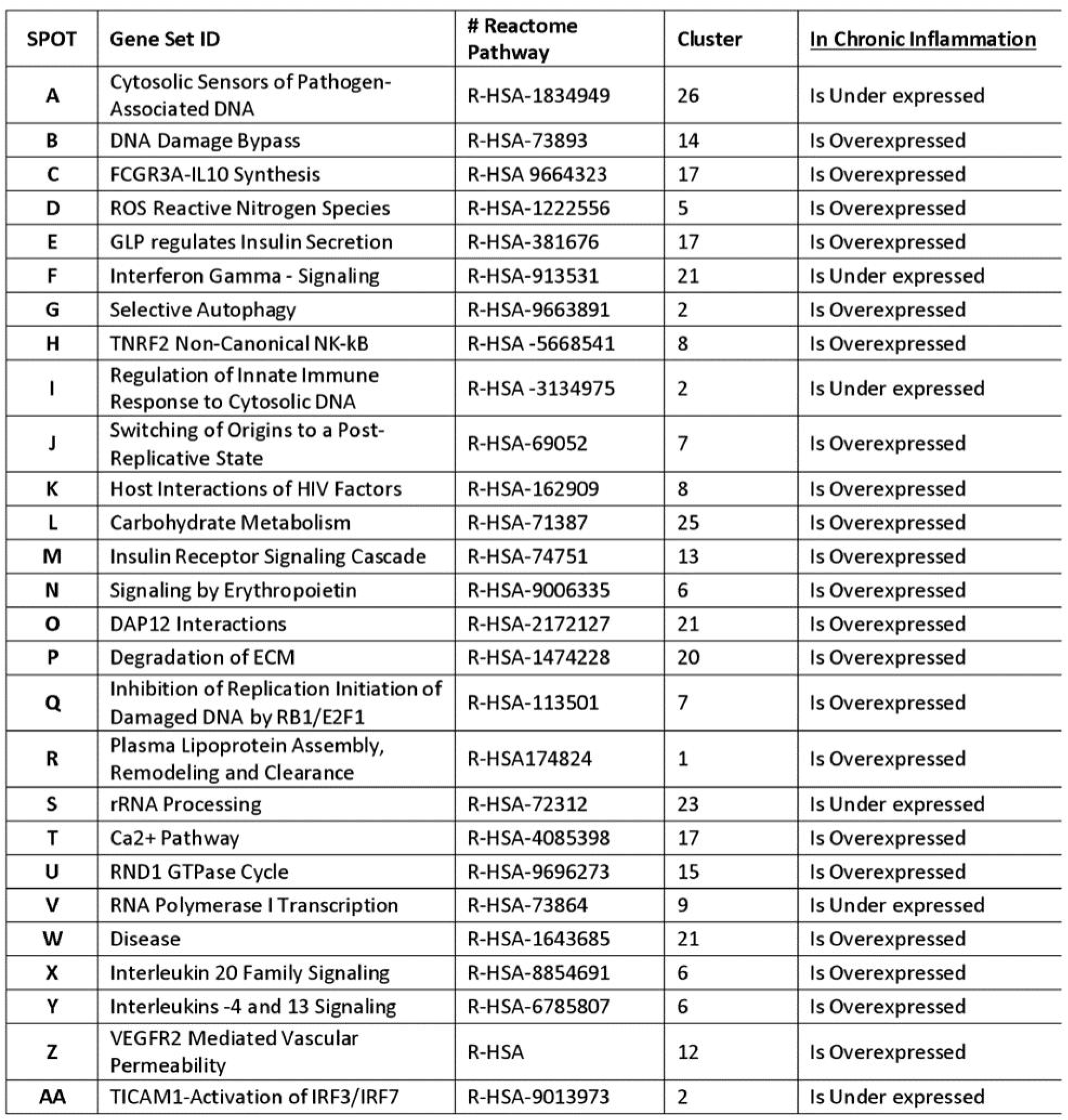
Tabulated Interpretation of Functional clustering as presented in Figure 3A. The data is defined by alphabetic value for spot, Library Gene set ID, annotated # of specific reactome pathways, cluster numbers and directional effect occurring in chronic inflammation.

A volcano plot generated from RNA-seq data uploaded via Reactome pathway bioinformatics revealed that chronic inflammation is associated with the overexpression of pathways presented in: **Figure 4 (top panel)** and the under expression of pathways presented **in Figure 4 (bottom panel)**. In brief, chronic inflammation leads to an increase in processes that regulate neutrophil degranulation, interleukin IL-4 and IL-13 signaling, RUNX keratinocytes and IL-10 and a reduction in processes that carry out antigen presentation, IFN signaling (extensive losses), C3 and C5 signaling and associated cytokines (CCL5), also presented as a full list of genes specific to functional ontology clustering in **Table 1 A** (Downregulated) and **Table 1 B** (Upregulated). The most notable overexpressed cyto/chemokines in chronic inflammation were the production of IL10 vs. resting [ +7.6 Log2FCp-Val 2.88E-14] and Ccl2 [6.0 Log2FC p-Val8.61E-19]. These genes were spot-tested for transcript‒protein correlations **(Figure 7 A, B, C).**

**Figure 4.**
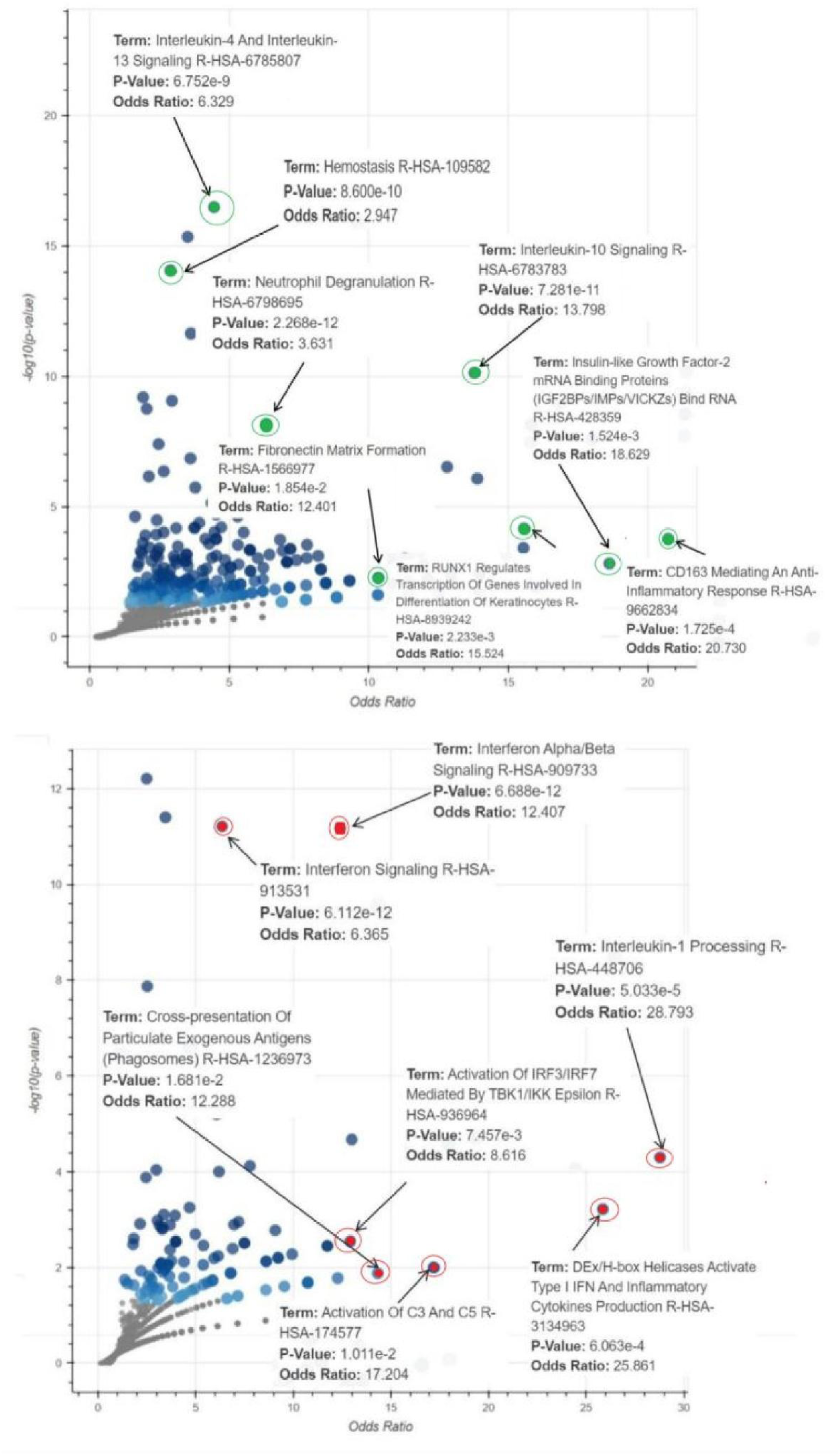
Functional clustering of events associated with chronic inflammation by Odds Ratio (Up regulated : Top Panel)/ (Down regulated :Bottom Panel). Functional biological events by odds ratio presented as volcano plots. The data was generated using analysis performed with the Reactome library. Each point represents a single Reactome gene set where the x-axis represents the odds ratio calculated for the gene set, whereas the y-axis represents the −log(p-value) of the gene set. The blue points represent significant terms associated with chronic inflammation (p<0.05); the gray points represent non-significant terms,the highlighted points [(red•)] (Bottom Panel) are for under expressed and (green•) (Top Panel) for over expressed in chronic inflammation with highest significance according to the p value and odds ratio. In-depth functional annotations are also provided in detail; Tables 1A (chronic under expressed) and 1B (chronic over expressed).

**Table 1 (A).**
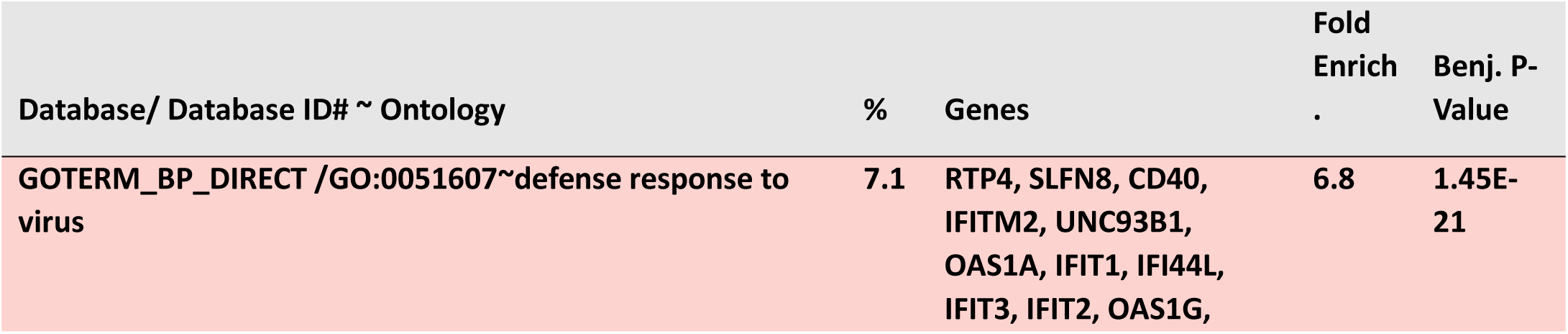
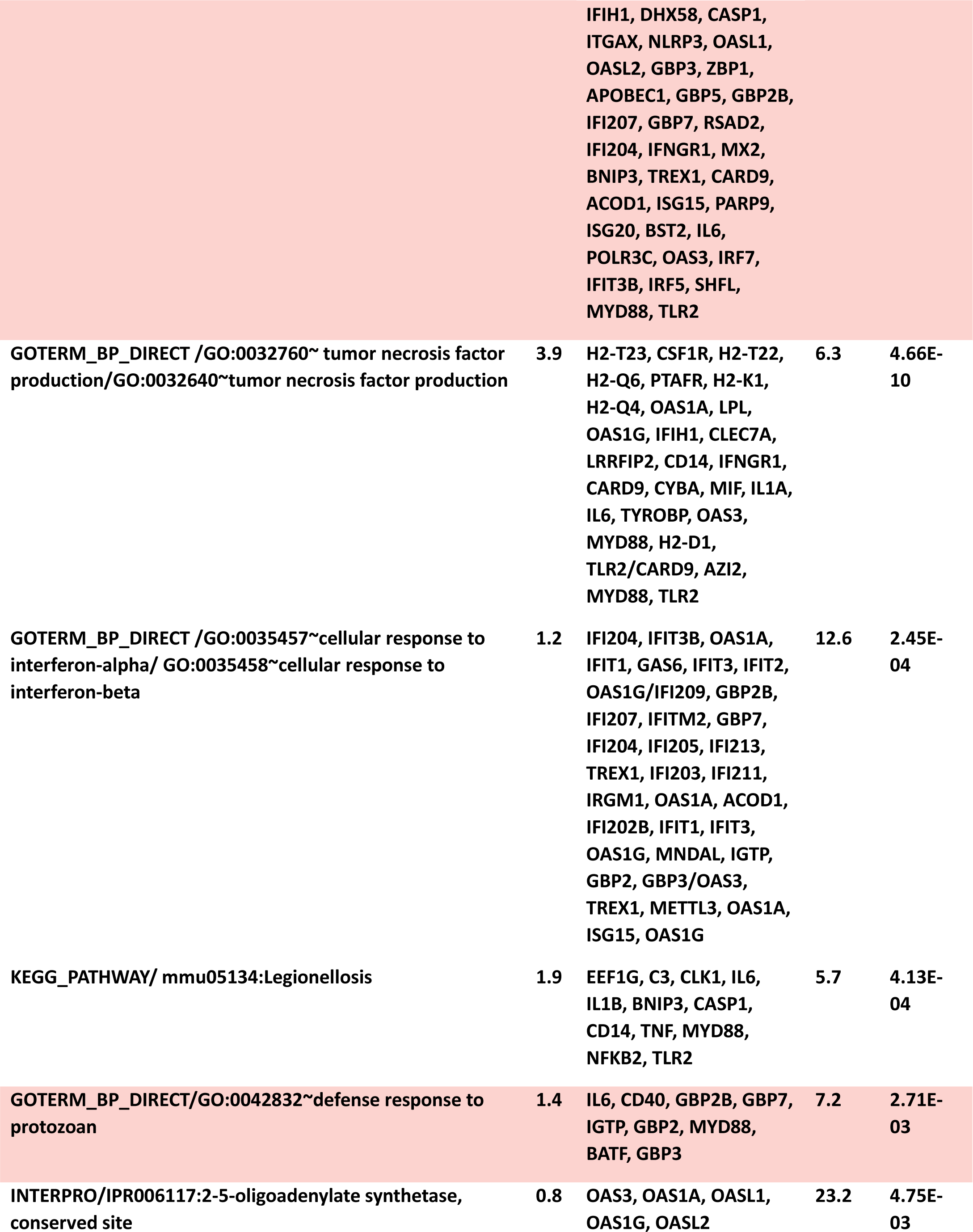
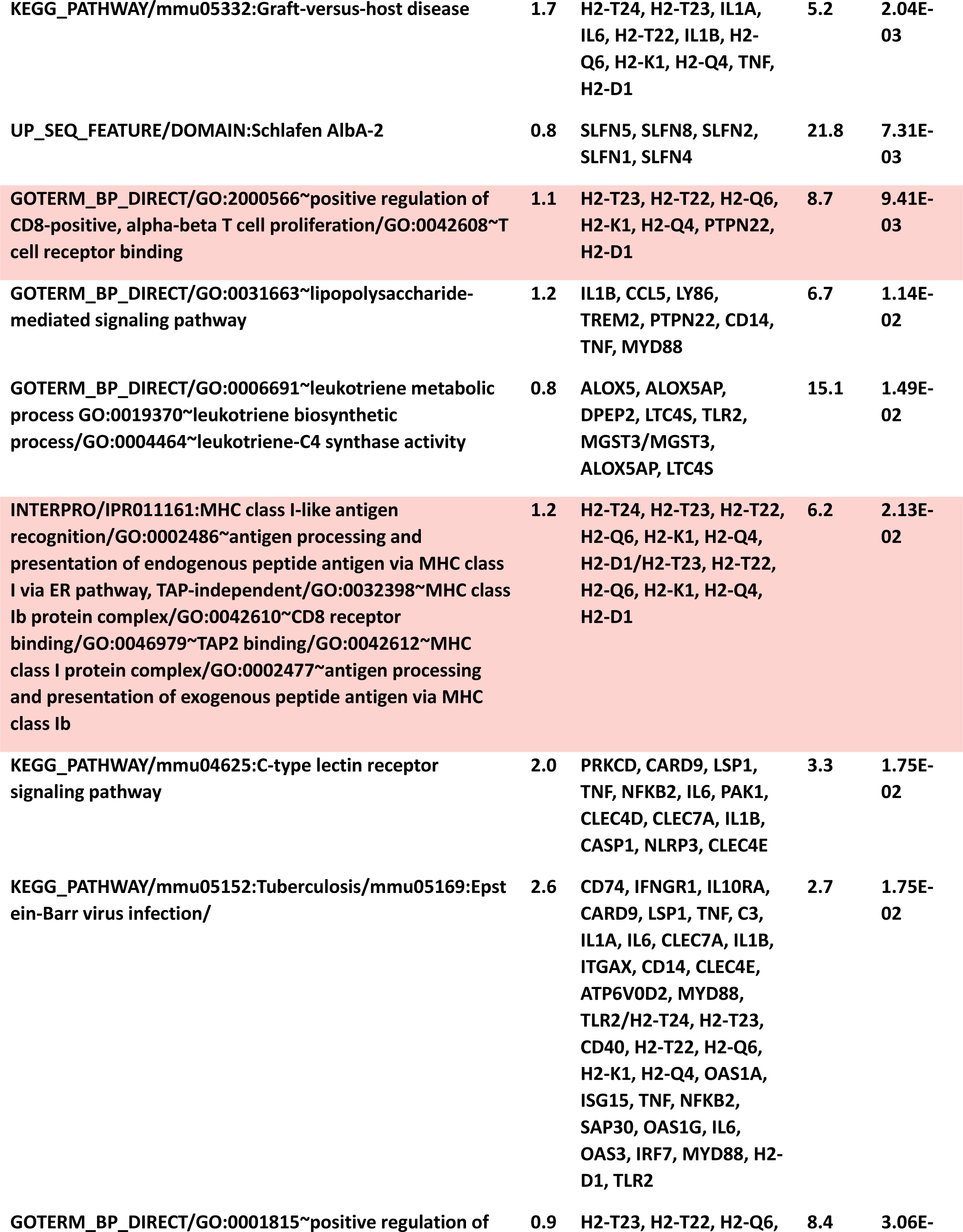
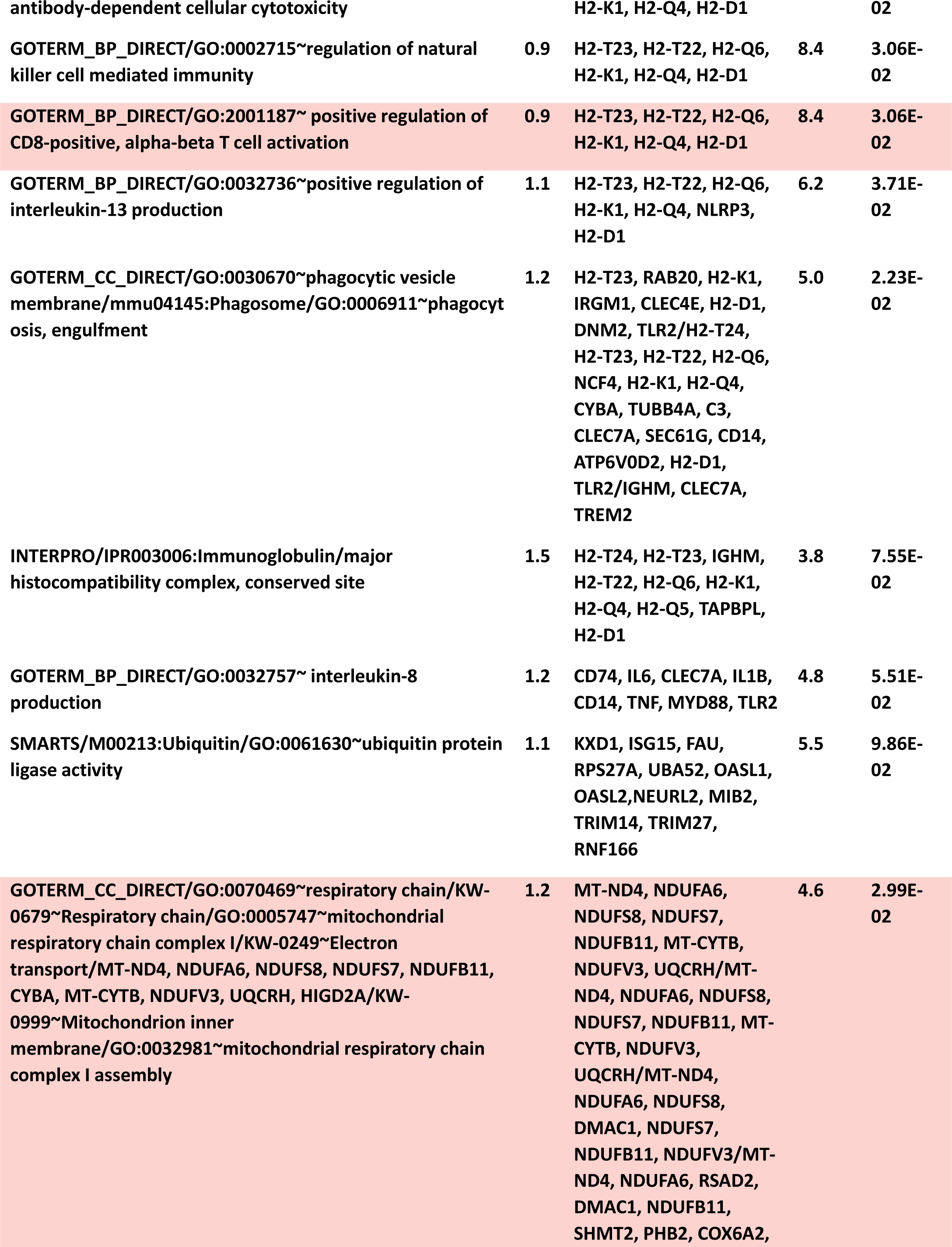
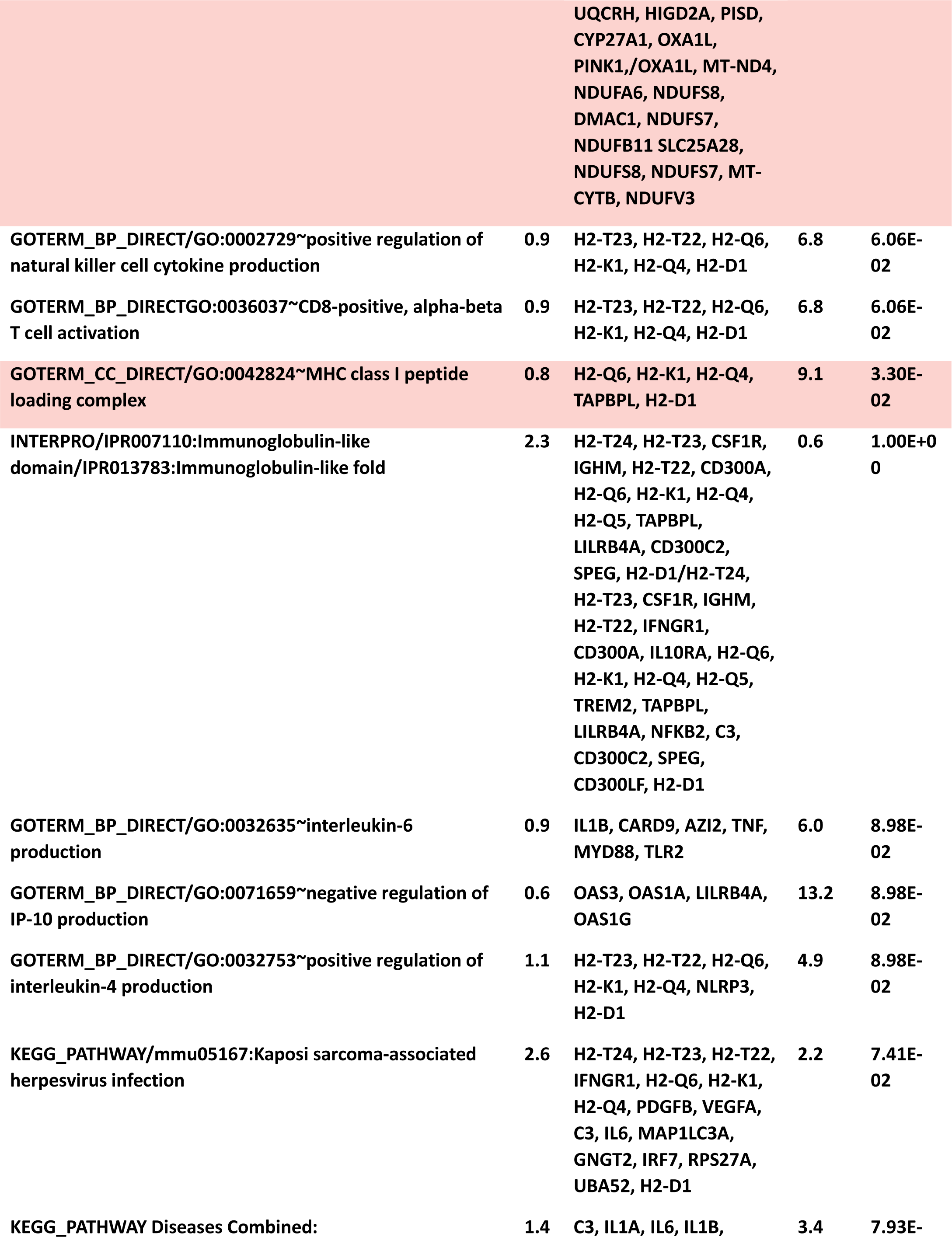
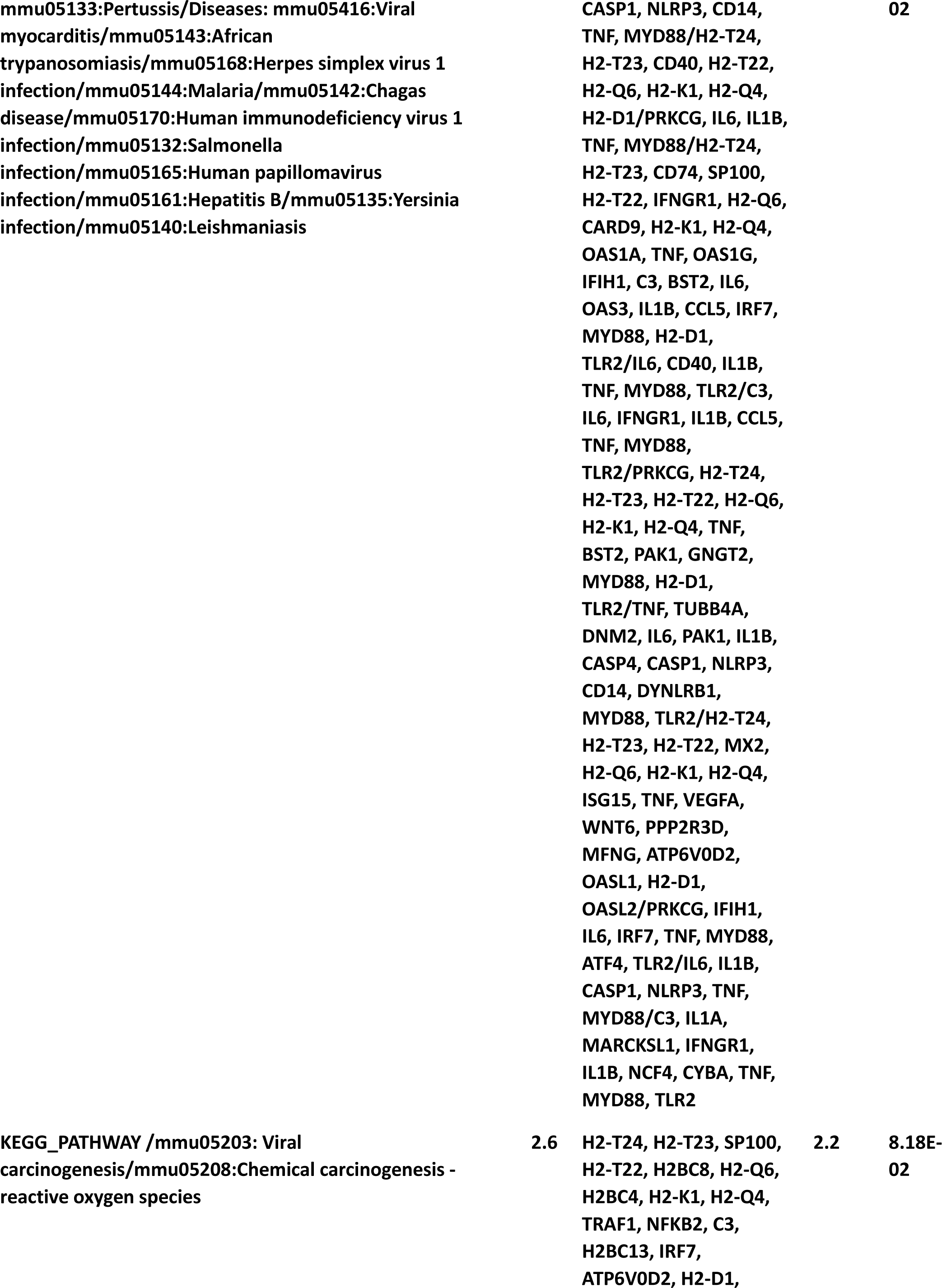
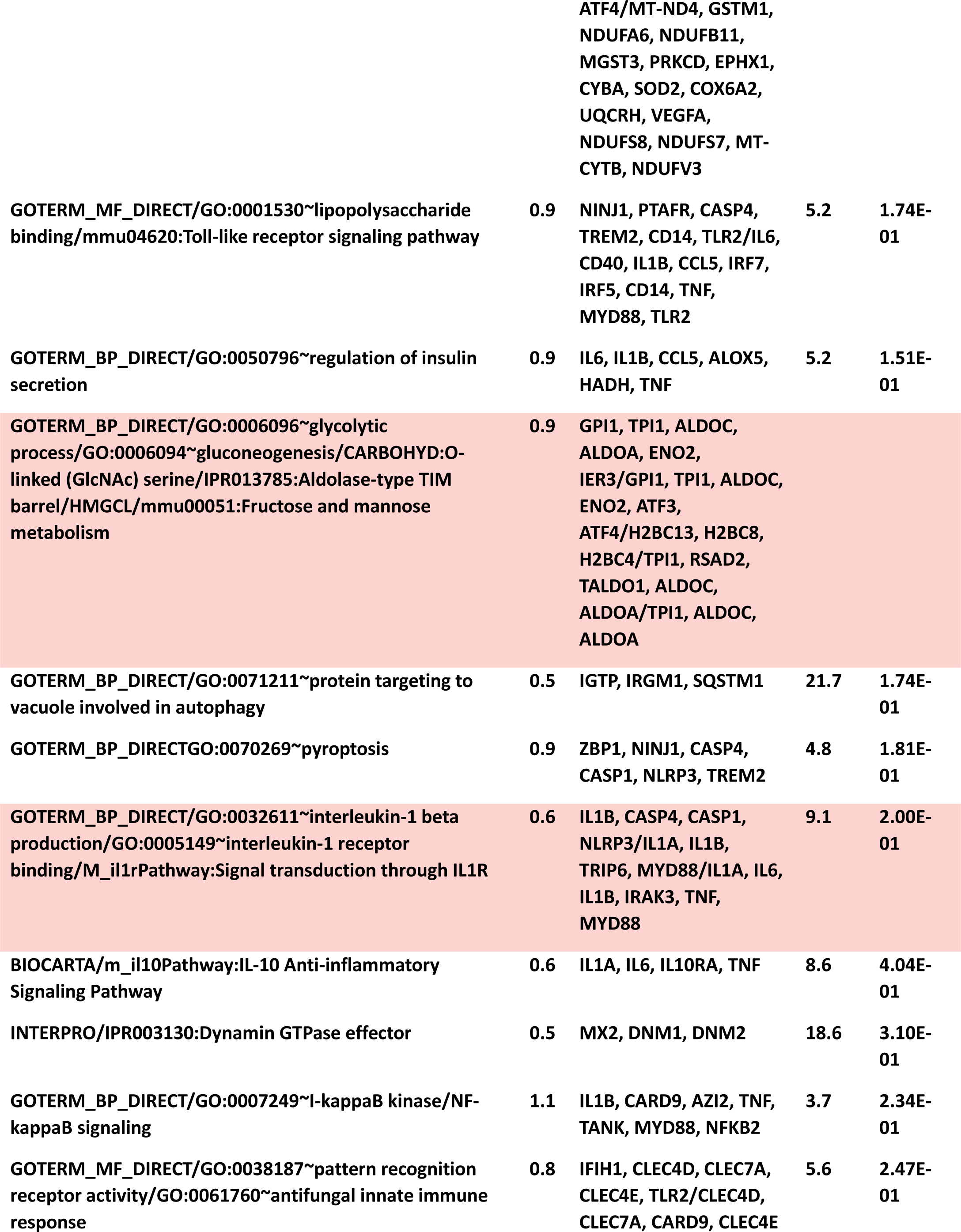
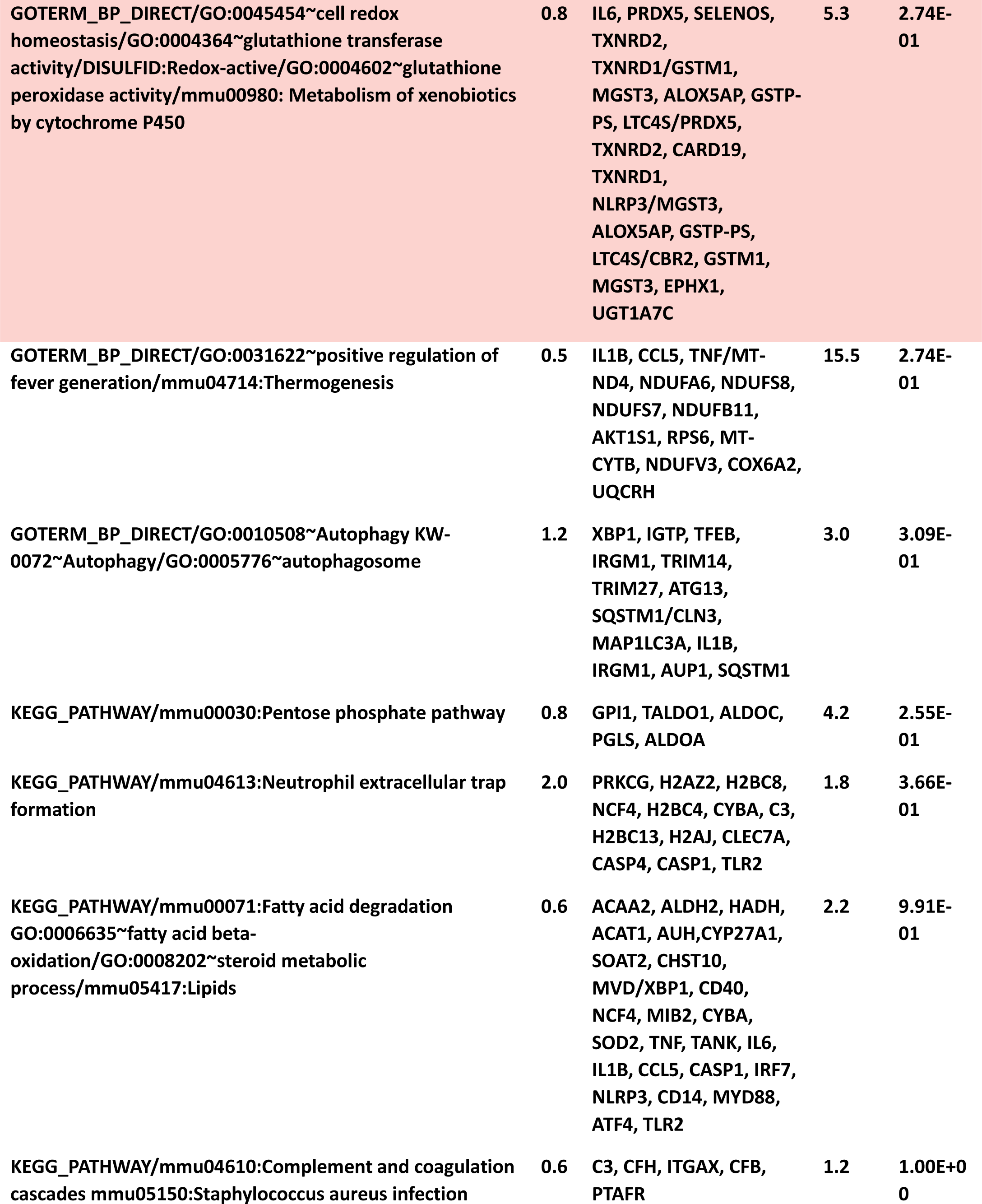
Downregulated.

**Table 1 B.** (Upregulated)

| Category /Term | % | Genes | Fold Enrich . | Genj P-Value |
| --- | --- | --- | --- | --- |
| GOTERM_CC_DIRECT/GO:0005764~lysosome/GO:0033176~proton-transporting V-type ATPase complex | 5.9 | ABCD4, CALCRL, SRC, SPPL2A, TCIRG1, GNS, CLN5, GJA1, SNX1, CTSL, CTSK, PSAP, CAPN2, ACP5, LRRC8A, SLC15A3, SIAE, CTSD, CTSC, SNX6, SRGN, CD164, SLC31A2, VPS33A, SLC11A1, ATP6AP2, FUCA2, SLC11A2, GGH, MT1, MCOLN2, SLC39A14, HCK, LMBRD1, MARCKS, RNF183, ATP6V0C, SQSTM1/ATP6V0C, ATP6V1D, ATP6V1C1 | 2.7 | 1.74E-05 |
| GOTERM_CC_DIRECT/GO:0001533~cornified envelope/GO:0030216~keratinocyte differentiation | 1.9 | SPRR2E, CSTA3, SPRR2F, SPRR2G, SERPINB2, SPRR2H, SPRR2I, SPRR2K, CSTDC4, SPRR2B, SPRR2D, STFA3/SPRR2E, CSTA3, SPRR2F, SPRR2G, SPRR2H, SPRR2I, SPRR2K, CFLAR, PTGS2, STFA3, CSTDC4, SPRR2B, SPRR2D | 6.1 | 3.03E-04 |
| INTERPRO /IPR000827:CC chemokine, conserved site/GO:0031726~CCR1 chemokine receptor binding/GO:0048247~lymphocyte chemotaxis/DOMAIN: Chemokine interleukin-8-like/KW-0145~Chemotaxis | 1.1 | CCL9, CCL12, CCL7, CCL6, CCL4, CCL3, CCL2/CCL9, CCL7, CCL6, CCL4, CCL3/CCL9, CCL12, CCL7, CCL6, CCL4, CCL3, CCL2, ADAM8//CCL9, CCL12, CCL7, CCL4, CCL3, CCL2, CXCL2/CCL9, CCL12, | 14.8 | 4.37E-03 |
|  |  | CCL7, CCL6, CCL4, C3AR1, CCL3, CCL2, FPR2, CXCL2, S100A8 |  |  |
| GOTERM_BP_DIRECT/GO:0071347~cellular response to interleukin 1/m_il22bpPathway:IL22 Soluble Receptor Signaling Pathway | 2 | CCL12, EDN1, SAA3, SERPINE1, CXCL2, CCL9, CCL7, CCL6, CCL4, ZC3H12A, CFL1, CCL3, CCL2/SOCS3, STAT1, JAK2, JAK1 | 5.3 | 2.18E-03 |
| GOTERM_BP_DIRECT/GO:0007264~small GTPase mediated signal transduction/GO:0043547~positive regulation of GTPase activity/IPR003578:Small GTPase superfamily, Rho type | 2.2 | DOCK5, DOCK4, VAV1, RND1, RHOB, RAP1B, DOCK11, RHOT2, RASGEF1B, HMOX1, RAPGEF2, RHOF, RAPGEF5, NUCB2/CCL12, RABGAP1L, ARHGAP35, VAV1, DOCK11, CCL9, CCL7, RASGEF1B, CCL6, RGS1, CCL4, RACK1, CCL3, CCL2, RAPGEF2, RAPGEF5, MAPRE2, MAP4K4/RHOT2, RHOF, RND1, RHOB | 4.3 | 4.98E-03 |
| UP_SEQ_FEATURE/DOMAIN:Ig-like V-type/KW-0393~Immunoglobulin domain | 2.2 | CD86, CD274, CD84, CD83, CD80, CEACAM1, SLAMF9, SLAMF7, CD300E, SIGLEC1, MPZL1, CD200, NECTIN2, HAVCR2/CD86, CD274, CD84, CD83, CD80, SEMA3E, LILRB4A, LY9, FCGR1, MILR1, CEACAM1, ALCAM, SLAMF9, SLAMF7, CD300E, SIGLEC1, FCGR2B, MPZL1, CD200, NECTIN2, MFAP3L, HAVCR2 | 4.2 | 1.15E-02 |
| GOTERM_BP_DIRECT/GO:0030593~neutrophil chemotaxis/GO:0048245~eosinophil | 1.9 | CCL9, CCL12, CCL7, SYK, CCL6, CCL4, SPP1, CCL3, CCL2, | 4.9 | 7.00E-03 |
| chemotaxis/GO:0002548~monocyte chemotaxis |  | CXCL2, VAV1, S100A8/CCL9, CCL12, CCL7, CCL6, CCL4, CCL3, CCL2/CCL9, CCL12, CCL7, CCL6, CCL4, CCL3, CCL2 |  |  |
| UP_SEQ_FEATUREDOMAIN:/Bcl-2 Bcl-2 homology region 1-3/GO:0008630~intrinsic apoptotic signaling pathway in response to DNA damage/GO:0097192~extrinsic apoptotic signaling pathway in absence of ligand/IPR026298:Bcl2 family/IPR020726:Apoptosis regulator, Bcl-2, BH2 motif, conserved site/IPR013282:Bcl-2-related protein A1 | 0.8 | BCL2L13, BCL2A1D, BCL2A1A, BCL2A1B, BCL2L1/PNP, MYC, EI24, HMOX1, BCL2A1D, TNFRSF1B, BCL2A1A, BCL2A1B, BCL2L1/LCN2, FAS, ITGAV, BCL2A1D, TNFRSF23, BCL2A1A, BCL2A1B, BCL2L1/BCL2L13, BCL2A1D, BCL2A1A, BCL2A1B, BCL2L1/BCL2A1D, BCL2A1A, BCL2A1B, BCL2L1/BCL2A1D, BCL2A1A, BCL2A1B | 19.7 | 2.17E-02 |
| UP_SEQ_FEATURE TNFR-Cys 3/IPR001368:TNFR/NGFR cysteine-rich region | 0.9 | FAS, TNFRSF10B, TNFRSF26, TNFRSF23, TNFRSF1B, TNFRSF22/FAS, TNFRSF10B, TNFRSF26, TNFRSF23, TNFRSF1B, TNFRSF22 | 11.2 | 3.86E-02 |
| GOTERM_BP_DIRECT/GO:0071356~cellular response to tumor necrosis factor | 2.3 | CCL12, EDN1, DHX9, FOXP1, NFKBIA, ZFP36, CCL9, FABP4, CCL7, CCL6, CCL4, ZC3H12A, CFL1, CCL3, CCL2 | 3.3 | 2.27E-02 |
| GOTERM_CC_DIRECT/GO:0005911~cell-cell junction/GO:0070161~anchoring junction | 2.9 | CD53, SH3KBP1, ACTN1, SLC2A1, ACTB, VAV1, RAP1B, AHI1, GJA1, CEACAM1, FLRT2, PDZD11, FABP7, CFL1, MICALL2, RAPGEF2, LCP2, ITGA5, NECTIN2/CD53, | 2.8 | 5.46E-03 |
|  |  | DOCK5, DBNL, TES, DST, SRC, SH3KBP1, ACTN1, PTPN12, RAP1B, SDCBP, AHI1, HCK, GJA1, CEACAM1, FLRT2, PDZD11, PDPN, MICALL2, RAPGEF2, ITGAV, ITGA5, HAVCR2 |  |  |
| GOTERM_BP_DIRECT/GO:0071346~cellular response to interferon-gamma/GO:0060333~interferon-gamma-mediated signaling pathway | 2 | CCL12, EDN1, NOS2, STAT1, CAPG, CCL9, CCL7, CCL6, CCL4, CCL3, CCL2, FLNB, CALM1/STAT1, JAK2, JAK1 | 3.7 | 2.48E-02 |
| INTERPRO/IPR004170:WWE domain | 0.8 | TIPARP, DTX2, ZC3HAV1, DTX4, PARP14 | 14.1 | 1.12E-01 |
| UP_KW_CELLULAR_COMPONENTKW-0458~Lysosome | 4.2 | ABCD4, SPPL2A, GNS, CLN5, CTSL, CTSK, PSAP, ACP5, LRRC8A, SLC15A3, SIAE, CTSD, CTSC, SRGN, CD164, SLC31A2, VPS33A, SLC11A1, ATP6AP2, SLC11A2, GGH, MCOLN2, SLC39A14, HCK, LMBRD1, RNF183, SQSTM1 | 2.2 | 4.42E-03 |
| KEGG_PATHWAY/mmu04062:Chemokine signaling pathway | 2.9 | CCR1, CCL12, SRC, STAT1, STAT2, PIK3CB, GNG12, CXCL2, VAV1, NFKBIA, RAP1B, HCK, CCL9, CCL7, CCL6, CCL4, CCL3, CCL2, JAK2 | 2.6 | 1.19E-02 |
| KEGG_PATHWAY/mmu04666:Fc gamma R-mediated phagocytosis | 1.9 | FCGR1, HCK, MARCKS, SYK, CFL1, PIP5K1B, PIK3CB, GAB2, FCGR2B, PLPP3, PLPP1, VAV1 | 3.4 | 1.88E-02 |
| UP_SEQ_FEATUREDOMAIN:SH3/GO:0042169~SH2 domain binding/DOMAIN:SH2/KW-0728~SH3 domain/KW-0727~SH2 | 2.5 | ARHGAP9, DOCK5, DOCK4, DBNL, DST, | 2.7 | 1.42E-01 |
| domain |  | SRC, NCF2, SH3KBP1, VAV1, AHI1, HCK, LASP1, UBASH3B, HCLS1, TSPOAP1, STAM2/SYK, SLC7A8, SRC, CTR9, RACK1, JAK2, SQSTM1/HCK, SOCS3, SYK, SRC, STAT1, STAT2, LCP2, JAK2, VAV1, JAK1/DOCK5, DOCK4, DBNL, DST, SRC, NCF2, SH3KBP1, VAV1, AHI1, HCK, LASP1, UBASH3B, HCLS1, TSPOAP1, STAM2/HCK, SOCS3, SYK, SRC, STAT1, STAT2, LCP2, JAK2, VAV1, JAK1 |  |  |
| GOTERM_MF_DIRECT/Metal Transporters GO:0005381~iron ion transmembrane transporter activity/GO:0034755~iron ion transmembrane transport/GO:0015086~cadmium ion transmembrane transporter activity/GO:0005384~manganese ion transmembrane transporter activity | 0.6 | TFRC, SLC11A1, SLC11A2, SLC39A14/SLC11A1, SLC11A2, LCN2, SLC39A14/SLC11A1, SLC11A2, SLC39A14/SLC11A1, SLC11A2, SLC39A14 | 16.3 | 9.94E-02 |
| GOTERM_BP_DIRECT/GO:0007259~JAK-STAT cascade/GO:0007167~enzyme linked receptor protein signaling pathway | 1.1 | CSF2RB2, SOCS3, STAT1, STAT2, CTR9, JAK2, JAK1/SYK, JAK2, JAK1 | 5.4 | 1.14E-01 |
| GOTERM_MF_DIRECT/GO:0035091~phosphatidylinositol binding/mmu04070:Phosphatidylinositol signaling system/GO:0046854~phosphatidylinositol phosphorylation/GO:0004438~phosphatidylinositol-3-phosphatase activity/mmu00562:Inositol phosphate metabolism | 1.7 | NISCH, SNX1, DAB2, HIP1, SNX12, ITPR2, TULP3, ITPR3, SNX10, STAM2, SNX6/ITPKB, PI4K2B, MTMR3, SYNJ1, PI4KA, ITPR2, ITPR3, PIP5K1B, PIK3CB, CALM1, IP6K2/ITPKB, PI4K2B, SOCS3, PI4KA, PIP5K1B, PIK3CB, IP6K2/ITPKB, PI4K2B, TPI1, MTMR3, SYNJ1, | 3.3 | 1.30E-01 |
|  |  | PI4KA, PIP5K1B, PIK3CB |  |  |
| KEGG_PATHWAY/mmu04064:NF-kappa B signaling pathway/IPR008343:Mitogen-activated protein (MAP) kinase phosphatase/GO:0033550~MAP kinase tyrosine phosphatase activity/GO:0043409~negative regulation of MAPK cascade | 1.9 | NFKBIA, SYK, GADD45A, CCL4, TRIM25, CFLAR, BCL2A1D, PTGS2, BCL2A1A, CXCL2, BCL2A1B, BCL2L1/DUSP4, DUSP5, DUSP1, DUSP16/DUSP4, DUSP5, DUSP1, DUSP16/SPRED3, DUSP4, DUSP5, DUSP1, RNF149, LILRB4A, DUSP16/MTMR10, MTMR3, SYNJ1, MTMR9 | 3 | 3.33E-02 |
| GOTERM_MF_DIRECT/GO:0005324~long-chain fatty acid transporter activity/GO:0015909~long-chain fatty acid transport/mmu00600:Sphingolipid metabolism/DOMAIN:Cytosolic fatty-acid binding proteins/GO:0042577~lipid phosphatase activity | 0.8 | ABCD4, FABP4, FABP5, SLC2A1, CD36/FABP4, FABP5, PLIN2, CD36/SGPL1, SPTLC2, PSAP, DEGS1, PLPP3, PLPP1, B4GALT5/FABP4, FABP5, FABP7/PLPP5, PLPP3, PLPP1 | 8.6 | 1.42E-01 |
| INTERPROIPR001452:Src homology-3 domain | 2.5 | ARHGAP9, DOCK5, DOCK4, DBNL, DST, SRC, NCF2, SH3KBP1, VAV1, AHI1, HCK, LASP1, UBASH3B, HCLS1, TSPOAP1, STAM2 | 2.4 | 3.14E-01 |
| GOTERM_MF_DIRECT/GO:0004715~non-membrane spanning protein tyrosine kinase activity/GO:0004725~protein tyrosine phosphatase activity | 1.1 | HCK, SYK, RIPK2, SRC, EIF2AK2, JAK2, JAK1/DUSP4, DUSP5, MTMR3, UBASH3B, DUSP1, PTPN12, SSH3, DUSP16, PTPN2 | 5 | 1.44E-01 |
| GOTERM_MF_DIRECT/GO:0005125~cytokine activity | 2.6 | IL10, IL33, CCL12, EDN1, IL1RN, CSF3, | 2.3 | 1.44E-01 |
|  |  | OSM, LIF, CXCL2, CCL9, CCL7, CCL6, CCL4, SPP1, CCL3, CCL2, TNFSF9 |  |  |
| INTERPRO/IPR001715:Calponin homology domain | 1.4 | SPECC1, DST, ACTN1, MICALL1, MICALL2, MICAL2, FLNB, MAPRE2, VAV1 | 3.7 | 3.14E-01 |
| KEGG_PATHWAY/mmu04216:Ferroptosis | 1.1 | TFRC, SLC11A2, VDAC3, HMOX1, SLC7A11, SAT1, SLC39A14 | 4.6 | 5.05E-02 |
| GOTERM_BP_DIRECT/GO:2001235~positive regulation of apoptotic signaling pathway/DOMAIN:PARP catalytic | 1.1 | CTSL, MYC, CTSK, OSM, FAS, JAK2, CTSC/PARP4, TIPARP, ZC3HAV1, PARP14 | 4.4 | 2.20E-01 |
| GOTERM_BP_DIRECT/GO:0002250~adaptive immune response | 2.9 | CD86, CD274, CD84, CLEC4A2, DBNL, SYK, CLEC4N, RIPK2, LILRB4A, MCOLN2, LY9, ADGRE1, CLEC4D, ALCAM, SLC7A8, SLAMF7, JAK2, HAVCR2, TRPM4 | 2.1 | 2.29E-01 |
| GOTERM_CC_DIRECT/GO:0030123~AP-3 adaptor complex | 0.6 | VPS33A, AP3D1, AP3B1, AP3S1 | 10.5 | 8.69E-02 |
| GOTERM_CC_DIRECT/GO:1904090~peptidase inhibitor complex/IPR000169:Cysteine peptidase, cysteine active site/IPR001713:Proteinase inhibitor I25A, stefin A | 0.6 | CSTA3, SERPINE1, CSTDC4, STFA3/CTSL, CTSK, CAPN2, CTSC/CSTA3, CSTDC4, STFA3 | 9.7 | 1.06E-01 |
| INTERPRO/IPR001433:Oxidoreductase FAD/NAD(P)-binding/KW-0285~Flavoprotein/IPR023173:NADPH-cytochrome p450 reductase, FAD-binding, alpha-helical domain-3/IPR008254:Flavodoxin/nitric oxide synthase | 0.6 | CYB5R1, POR, MTRR, NOS2/CYB5R1, POR, MTRR, NOS2, SMOX, MAOA, TXNRD1, MTHFR, MICAL2, QSOX1, PPCDC/POR, MTRR, NOS2/POR, MTRR, NOS2 | 9.7 | 5.01E-01 |
| UP_KW_BIOLOGICAL_PROCESS/KW-0395~Inflammatory response | 2.3 | CCL12, DHX9, CXCL2, NDST1, ATRN, HCK, | 2.2 | 9.14E-02 |
|  |  | CCL7, TNIP1, CCL4, ZC3H12A, STAB1, CCL3, CCL2, S100A8, HAVCR2 |  |  |
| GOTERM_BP_DIRECT/GO:0006511~ubiquitin-dependent protein catabolic process/GO:0016567~protein ubiquitination | 2.6 | AREL1, PSMD11, USP32, USP12, UBE2D3, UBE4A, USP18, BAG6, PSMA6, HERC3, RNF149, PSMC2, RLIM, ARIH2, TRIM25, HERC6, UBL7/TRIM12C, TRIM30A, AREL1, MGRN1, UBE2D3, MED8, UBE4A, DTX2, DTX4, MED7, RNF41, SOCS3, HERC3, UBR5, SDE2, TRIM13, RACK1, RLIM, ARIH2, RNF183, FAU, HERC6 | 2 | 3.89E-01 |
| UP_KW_BIOLOGICAL_PROCESS/KW-1064~Adaptive immunity | 2.9 | CD86, CD274, CD84, CLEC4A2, DBNL, SYK, CLEC4N, RIPK2, LILRB4A, MCOLN2, LY9, ADGRE1, CLEC4D, ALCAM, SLC7A8, SLAMF7, JAK2, HAVCR2, TRPM4 | 1.9 | 1.26E-01 |
| INTERPRO/IPR011993:Pleckstrin homology-like domain | 3.4 | ARHGAP9, OSBPL8, MTMR10, RABGAP1L, APLP2, PLEK, RAPH1, MSN, MTMR9, TBC1D9, GAB2, RASAL2, VAV1, PLEKHA8, SPRED3, DOCK11, DAB2, FGD6, RASA2, WDFY3, PLEKHN1, JAK2 | 1.8 | 0.766 |
| GOTERM_BP_DIRECT/GO:0046777~protein autophosphorylation | 2 | SYK, SRC, NEK6, EIF2AK2, IRAK3, STK10, HCK, STK17B, PIM1, RIOK2, PIM2, | 2.2 | 4.46E-01 |
|  |  | JAK2, CALM1 |  |  |
| GOTERM_BP_DIRECT/GO:0035654~cargo loading into clathrin-coated vesicle, AP-3-mediated | 0.5 | AP3D1, AP3B1, AP3S1 | 14.2 | 4.66E-01 |
| GOTERM_BP_DIRECT/GO:0045087~innate immune response | 5.3 | TRIM12C, CD84, CLEC4A2, TRIM30A, DHX9, SRC, CLEC4N, ZDHHC5, ZC3HAV1, LY9, NPY, CLEC5A, SLAMF7, TRIM25, WFDC17, JAK2, SLC15A3, HERC6, UBL7, HAVCR2, SYK, XRCC5, RIPK2, SP110, RIOK3, EIF2AK2, PARP14, MCOLN2, FCGR1, HCK, CLEC4D, TRIM13, LCN2, S100A8 | 1.5 | 4.79E-01 |
| UP_SEQ_FEATUREREGION/:Phosphatase sequence motif I/GO:0004721~phosphoprotein phosphatase activity/IPR003595:Protein-tyrosine phosphatase, catalytic | 0.5 | PLPP5, PLPP3, PLPP1/DUSP4, DUSP5, MTMR3, UBASH3B, DUSP1, PPM1H, PTPN12, SSH3, DUSP16, PTPN2/DUSP4, DUSP5, MTMR3, DUSP1, PTPN12, PTPN2 | 13.3 | 9.19E-01 |
| UP_SEQ_FEATUREDOMAIN/:bMERB/IPR022735:Domain of unknown function DUF3585 | 0.5 | MICALL1, MICALL2, MICAL2 | 13.3 | 9.19E-01 |
| INTERPRO/IPR016186:C-type lectin-like | 1.6 | ATRN, CLEC4D, CD72, CLEC4A2, CLEC4N, KLRA2, CLEC5A, STAB1, CD69, CD44 | 2.5 | 8.22E-01 |
| GOTERM_BP_DIRECT/GO:0042130~negative regulation of T cell proliferation | 0.9 | CD86, CD274, CEACAM1, CD80, LILRB4A, HAVCR2 | 3.7 | 5.52E-01 |
| GOTERM_BP_DIRECT/GO:0060155~platelet dense granule organization/IPR003531:Short hematopoietin receptor, family 1, conserved site | 0.6 | AP3D1, AP3B1, AP3S1, AP1S3/CSF2RB2, IL4RA, IL7R | 5.8 | 6.38E-01 |
| GOTERM_BP_DIRECT/GO:0046596~regulation of viral entry | 0.6 | TRIM12C, TRIM30A, | 5.8 | 6.38E-01 |
| into host cell/GO:0044790~negative regulation by host of viral release from host cell |  | TRIM25, NECTIN2/TRIM12C, TRIM30A, TRIM13, TRIM25 |  |  |
| INTERPRO/IPR013766:Thioredoxin domain | 0.8 | TXNDC9, PRDX1, QSOX1, DNAJC16, PDIA4 | 4.1 | 9.96E-01 |
| INTERPRO/IPR026791:Dedicator of cytokinesis | 0.5 | DOCK11, DOCK5, DOCK4 | 9.3 | 1.00E+00 |
| GOTERM_BP_DIRECT/GO:0030574~collagen catabolic process/GO:0001968~fibronectin binding | 0.8 | CTSL, CTSK, MMP19, MMP8, MMP9/CTSL, CTSK, ITGAV, LILRB4A, MMP9 | 3.5 | 8.34E-01 |
| UP_KW_BIOLOGICAL_PROCESSK/W-0644~Prostaglandin metabolism | 0.5 | PTGS2, PTGR1, PTGES | 6.8 | 5.04E-01 |
| KEGG_PATHWAY/mmu05235:PD-L1 expression and PD-1 checkpoint pathway in cancer | 0.9 | NFKBIA, CD274, STAT1, PIK3CB, JAK2, JAK1 | 1.8 | 8.12E-01 |

Using the DAVID function ontology, KEGG pathways were generated for greater specificity with respect to the pathway affected and genes under that pathway in the chronic gene sets presented in **Figure 5** (Jak/Stat-IL-10) and **Figure 6** (cytokine–cytokine–receptor interaction), with the legend listing DEGs by Log2FC, p value. KEGG diagrams reflect upregulated transcripts by box highlighting, with the text of the gene symbol and designated with a [+], except for a single loss of CCL4 atypical of the entire series of chemokines; designated with a red box [-]. Moreover, the results of the KEGG analysis of the chronic-inflammatory-urendersexpd DEGs are presented in **Figure 8**, which shows the losses of transcripts required for antigen presentation/major histocompatibility: H2T23, H2-T22, H2-Q6, H2-K1, H2-Q4, PTPN22, H2-D1 and dozens of transcripts that are required for host type I IFN signaling, as presented in Table 1 A (IFI204, IFIT3B, OAS1A, IFIT1, GAS6, IFIT3, IFIT2, OAS1G/IFI209, GBP2B, IFI207, IFITM2, GBP7, IFI204, IFI205, IFI213, TREX1, IFI203, IFI211, IRGM1, OAS1A, ACOD1, IFI202B, IFIT1, IFIT3, OAS1G, MNDAL, IGTP, GBP2, GBP3/OAS3, TREX1, METTL3, OAS1A, ISG15, and OAS1G). Moreover, as shown in **Figure 9**, chronic inflammation leads to major losses in NOD signaling, creating defects in defense against pathogenic bacteria, RNA viruses, and responses to PAMPs, DAMPs, and bacterial proteoglycans.

**Figure 5.**
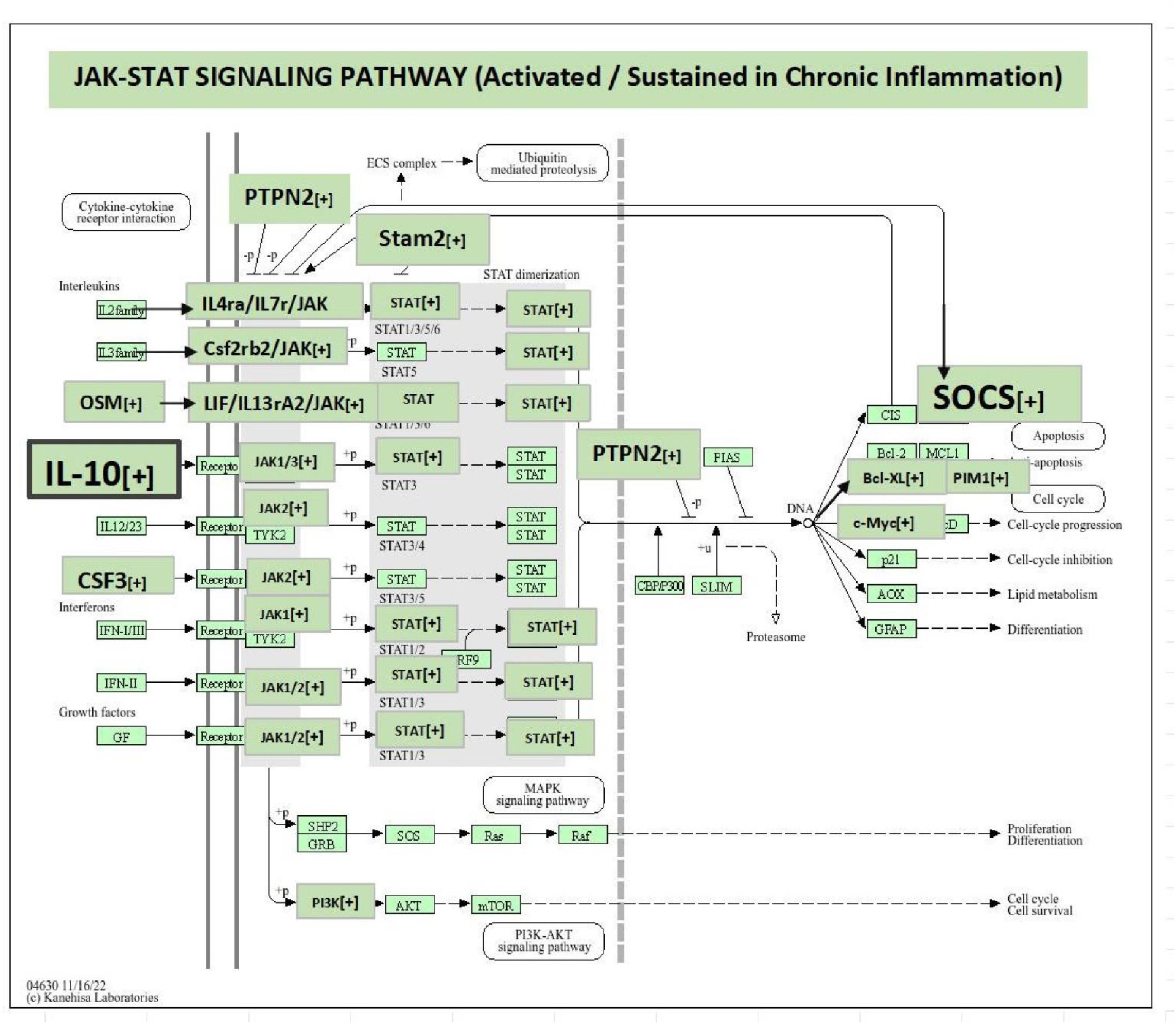
Upregulated in Chronic; Jak-Stat: IL-10 Pathway: DEGs up regulated in chronic inflammation were pooled into DAVID pathway analysis with pathway analysis showing significant highlighting changes in the Jak/Stat pathway. Major changes are presented by Green overlay text boxes with noted changes in Gene, FC and p-value for; **Il10** [+7.6 Log2FC,p-Val =2.88E-14], **\*Socs3** [+4.9 Log2FC,p-Val 8.73E-20], **Lif** [+3.7Log2FC,p-Val =3.57E-08], **Pim2** [+ 3.0Log2FC,p-Val =2.31E-06],**\*Csf3** [+3.4 Log2FC, p-Val =4.55E-02] and **Ptpn2** [+2.9 Log2FC, p-Val =1.53E-04].

**Figure 6.**
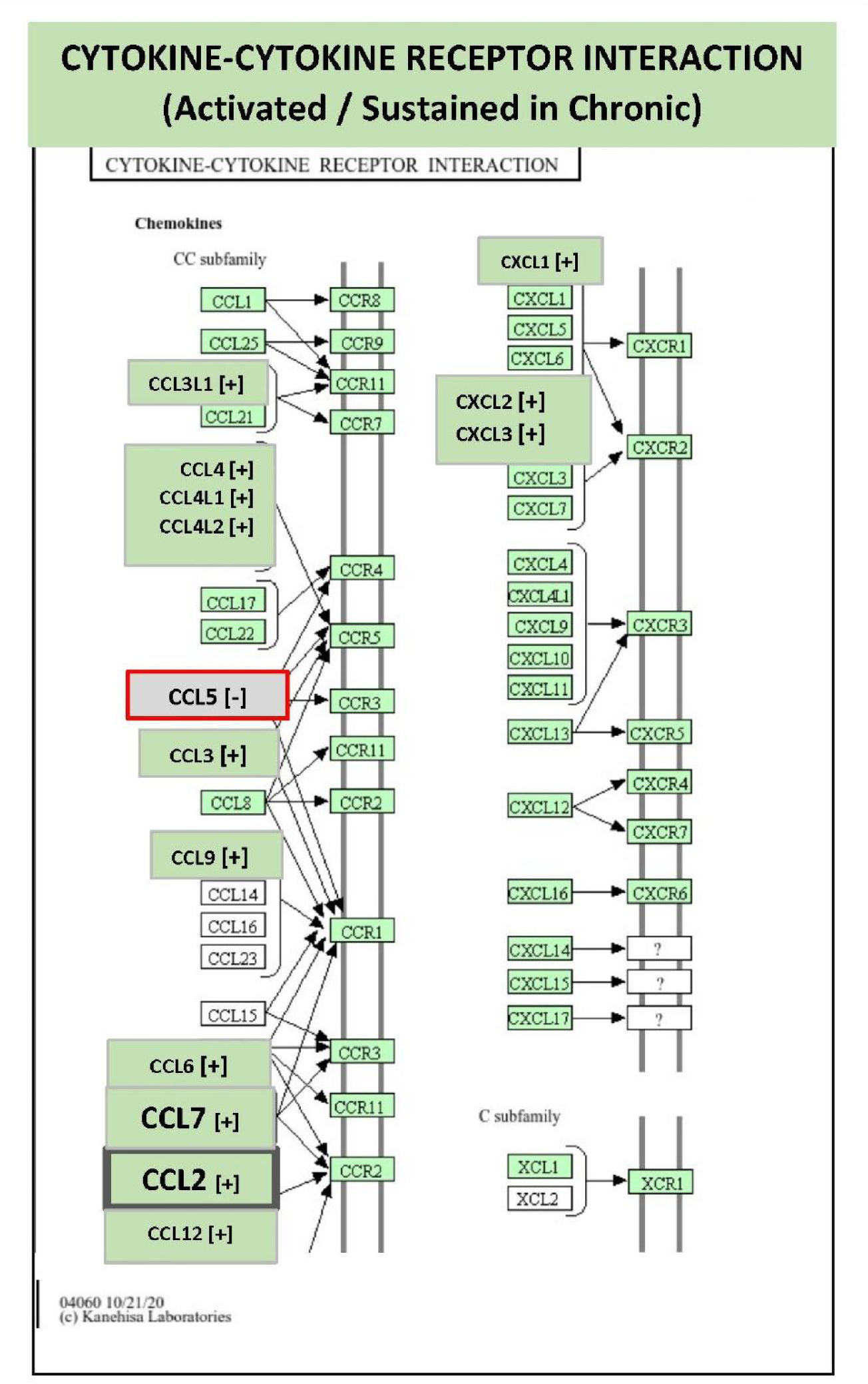
Upregulated in Chronic; Cytokine −Cytokine Receptor Interaction: CCL2/CCL7 Signaling. Upregulated DEGs associated with chronic inflammation were pooled into DAVID pathway analysis, which revealed significant impact on the cytokine/cytokine interreceptor interaction pathway. Major changes by gene, FC and p-value were found for : **\*CCL7** [ +5.8 Log2FC,p-Val 1.79E-06], **\*Ccl2** [+ 6.0 Log2FC, p-Val 8.61E-19] and **\*CL12** [+5.8 Log2FC,p-Val 1.36E-03]. There was singular significant loss in CCL 5[-].

**Figure 7A.**
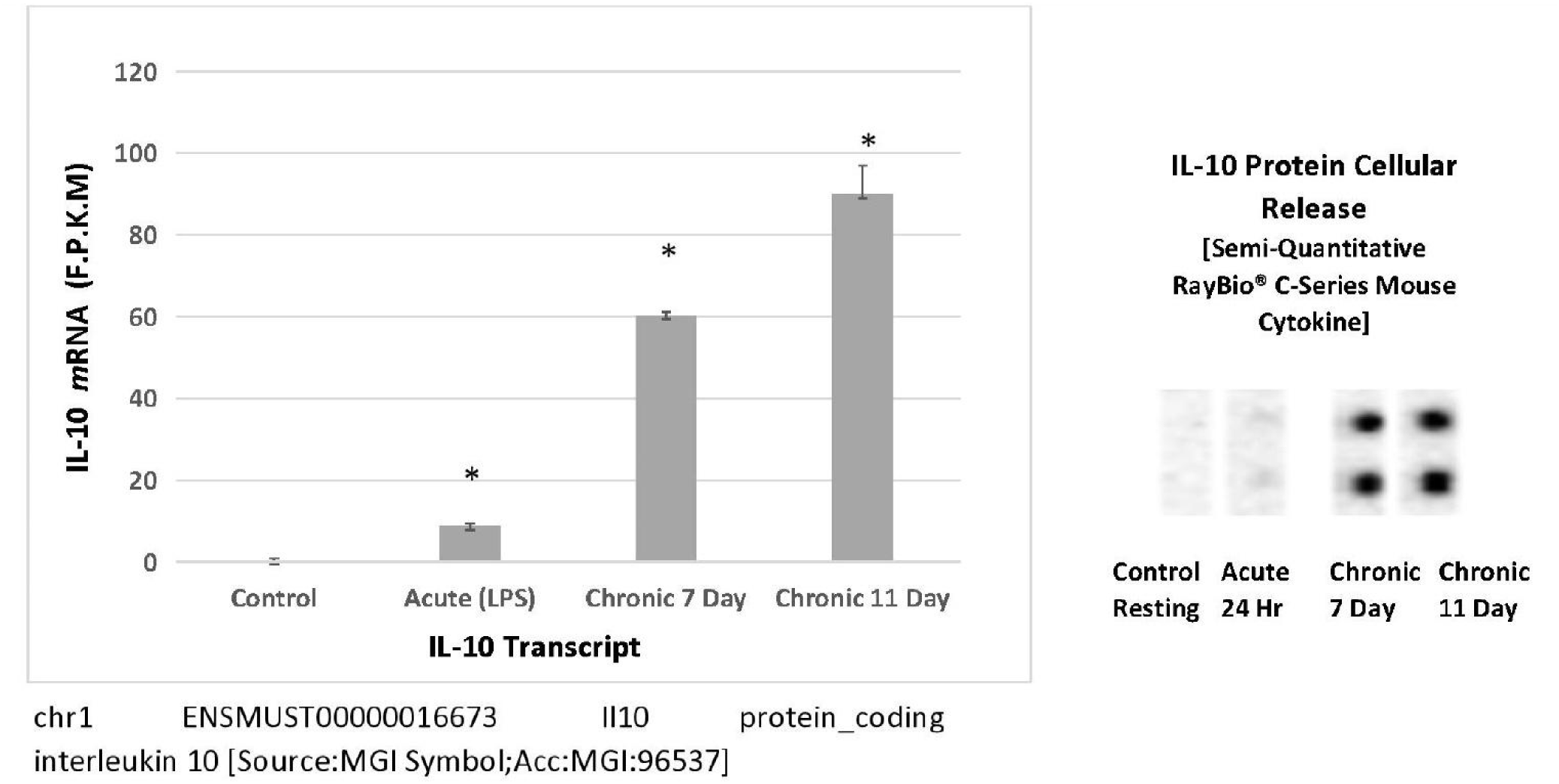
Transcription to Protein Translation Alignment for: IL-10. IL-10 was evaluated by RNA-Seq and an antibody array for the detection of proteins released into the cell supernate. The data are expressed as the mean ± S.E.M FPKM= [total_exon_fragments /mapped_reads (millions) × exon length(kB)]), n=3, significance from controls *p<0.001 and densitometry spots as imaged by chemiluminescence on a Bio-Rad Versa doc imager.

**Figure 7B.**
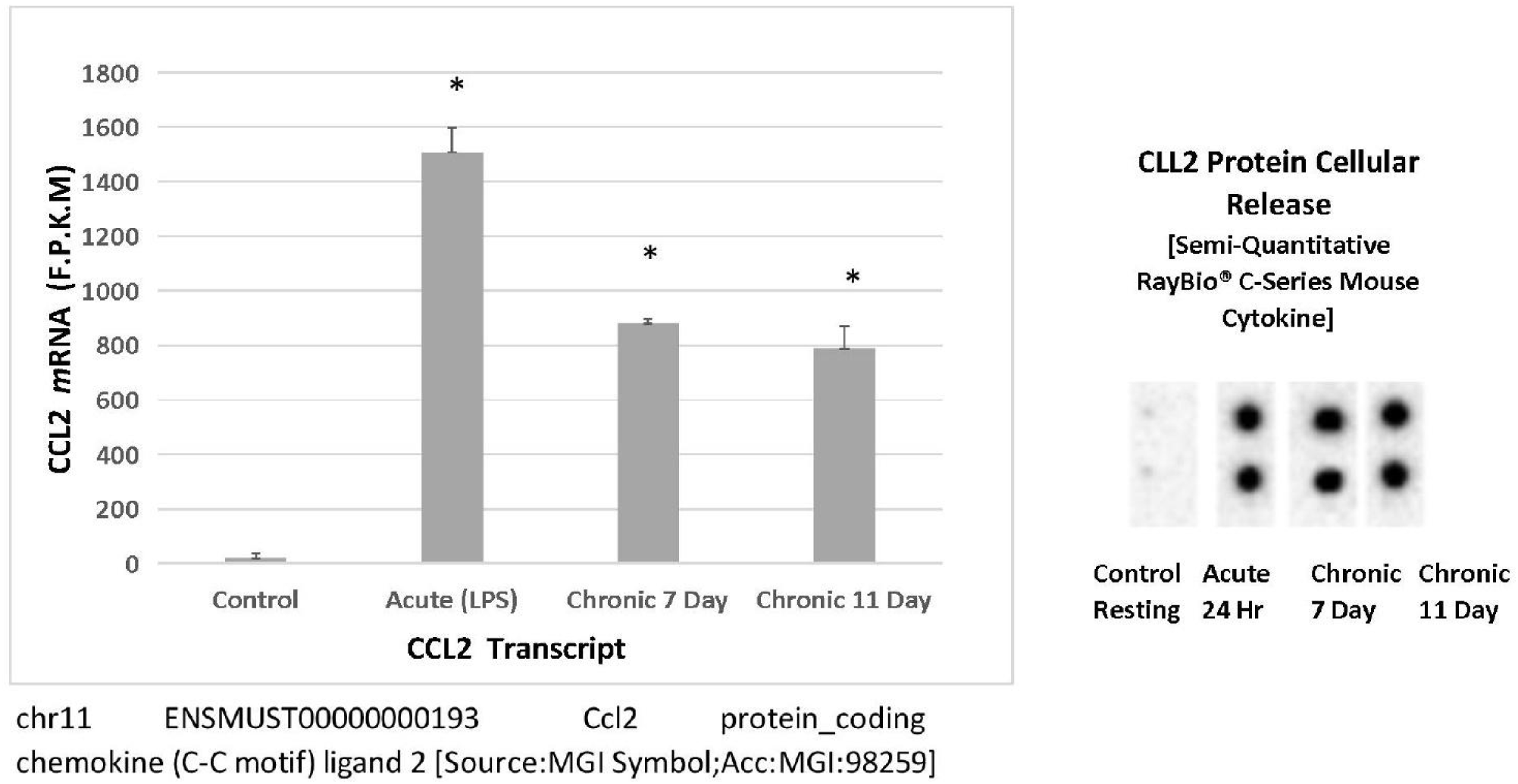
Transcription to Protein Translation Alignment; chemokine CCL2. CCL2 was evaluated by RNA-Seq and an antibody array for the detection of proteins released into the cell supernate. The data are expressed as the mean ± S.E.M FPKM= [total_exon_fragments /mapped_reads (millions) × exon length(kB)]), n=3, significance from controls *p<0.001 and densitometry spots as imaged by chemiluminescence on a Bio-Rad Versa doc imager.

**Figure 7C.**
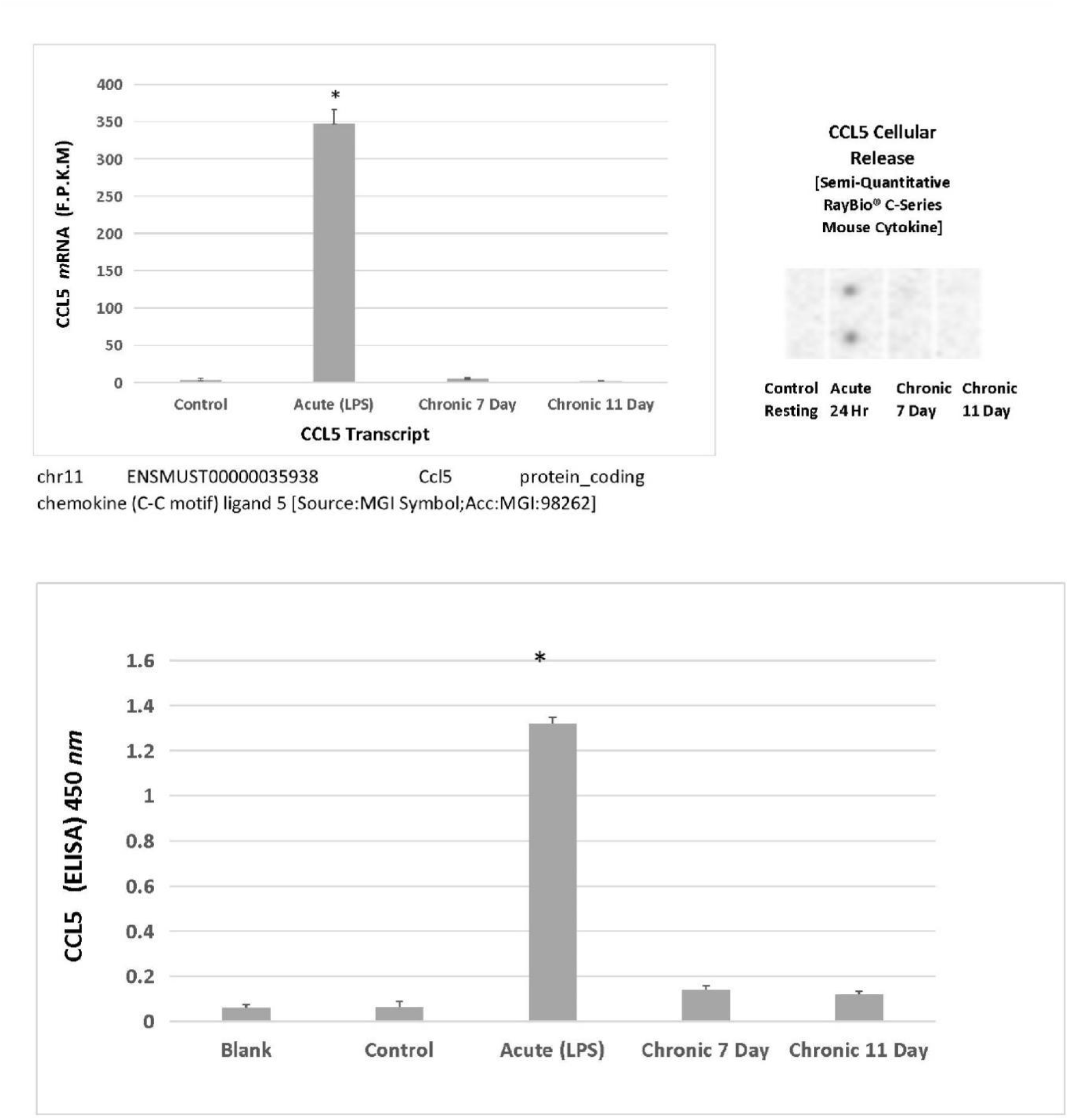
Transcription to Protein Translation Alignment; chemokine CCL5. CCL5 was evaluated by RNA-Seq an antibody array and an ELISA for the detection of protein released into the cell supernate. The data are expressed as the mean ± S.E.M FPKM= [total_exon_fragments /mapped_reads (millions) × exon length(kB)]), n=3, significance from controls *p<0.001 and densitometry spots as imaged by chemiluminescence on a Bio-Rad Versa doc imager. **Figure 7D.** ELISA confirmation of CCL5 protein release into the cell supernatant. The data represent the O.D. at 450 nm and are expressed as the mean ± S.E.M., n=3. A significant difference from the control was determined via one-way ANOVA, followed by a Tukey post hoc test. *p<0.05.

**Figure 8.**
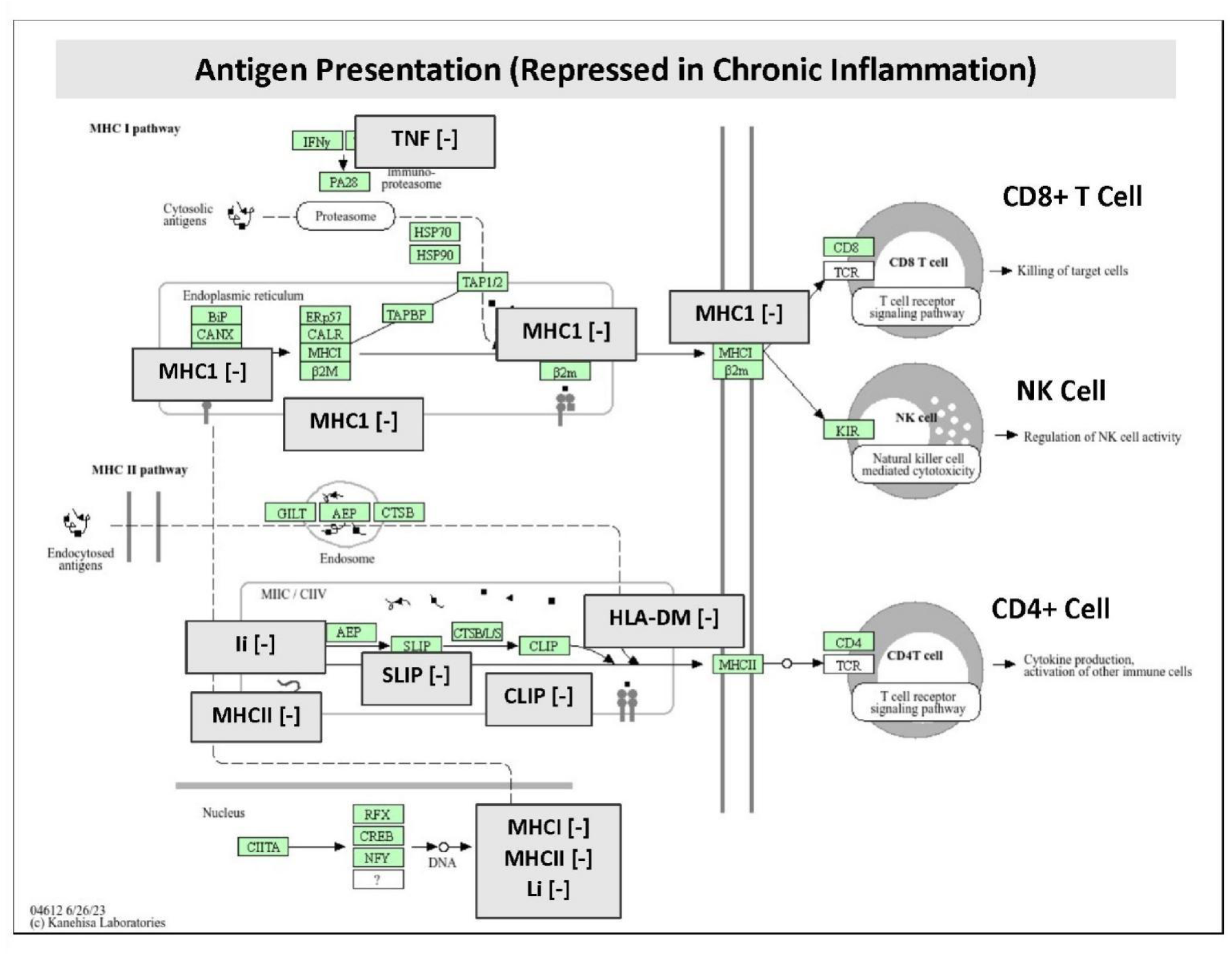
Downregulated in Chronic; MHC1 MCH2 Antigen Presentation. Downregulated DEGs associated with chronic inflammation were pooled into DAVID pathway analysis, which revealed significant impact on the antigen presentation. Changes are reflected by text overlay boxes and significant losses were found for; Gene, FC, p-Val; **MCH1;** (**\*H2-T24** −2.60 Log2FCp-Val=1.66E-73),(**\*H2-M2** −4.87 Log2FC p-Val=2.72E-53),(**\*H2-T23** −3.10 Log2FC p-Val=6.45E-47),(**\*H2-T10** −2.78, Log2FC p-Val = 2.22E-14),(**\*H2-K2** −2.01 Log2FCp-Val=2.86E-12), (**\*H2-M2** −6.50 Log2FC p-Val= 3.72E-12), (**\*H2-K1** −17.02 Log2FC p-Val=5.57E-10),(**\*H2-T3** −9.97 Log2FC p-Val =8.01E-05),(**\*H2-K1** −3.98 Log2FC p-Val=1.21E-04),(**\*H2-T22** −2.29 Log2FC p-Val=2.36E-04),(**\*H2-Q2** −10.87 Log2FCp-Val=5.75E-04),(**\*H2-Eb1** −9.74 Log2FC p-Val = 8.20E-04),(**\* H2-Q5** −18.53 Log2FC p-Val = 1.78E-03),(**\*H2-Q1** −15.89 Log2FCp-Val=5.71E-03),(**Li, Slip and Clip (CD74)** −3.4 Log2FC p-Val=2.23E-06).

**Figure 9.**
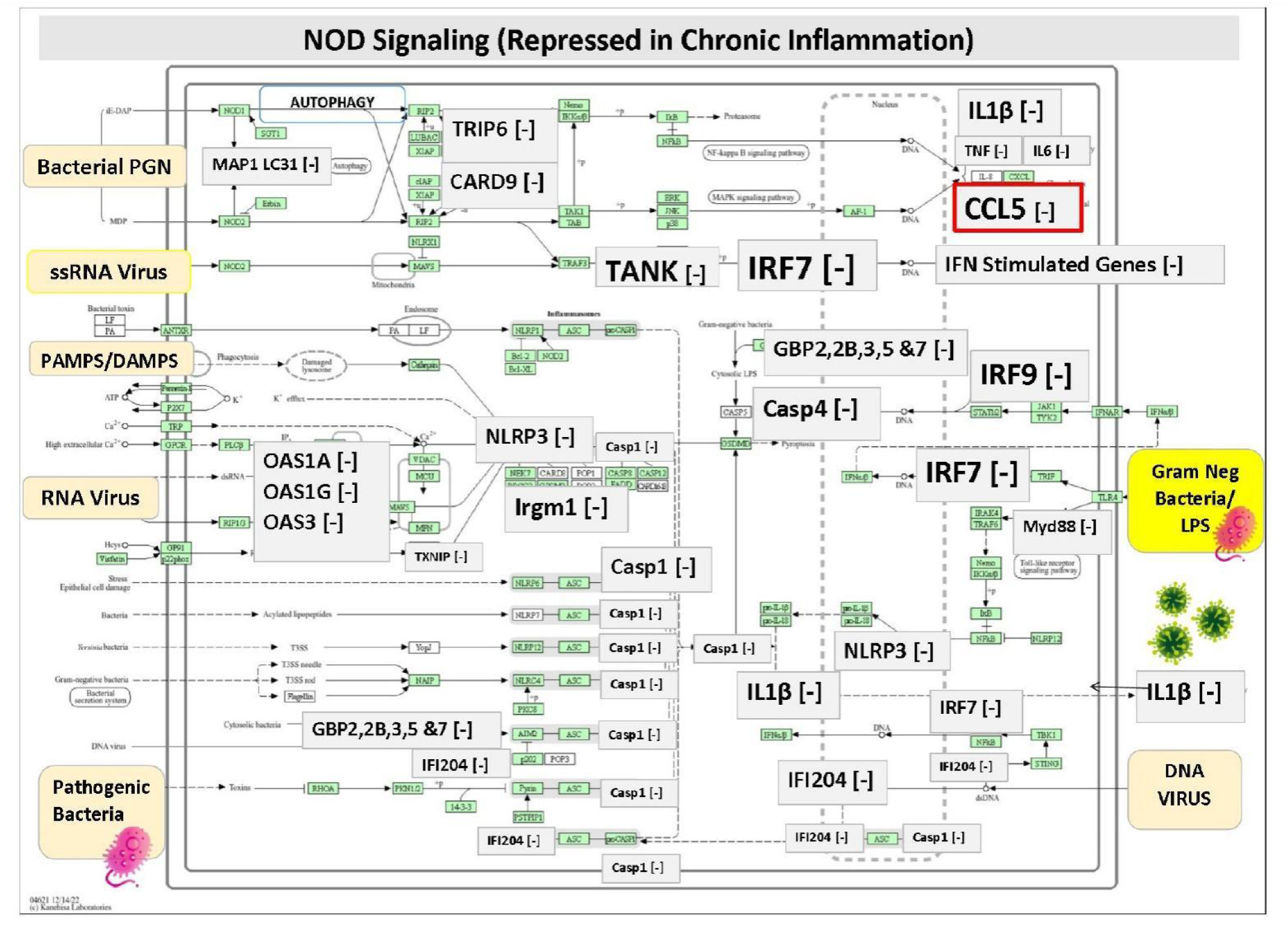
Downregulated in Chronic; NOD Signaling. Downregulated DEGs associated with chronic inflammation were pooled into DAVID pathway analysis, which revealed significant negative impact on NOD Signaling. Changes are reflected by text overlay boxes and significant losses were found for; Gene, FC, p-Val; (**\*Nlrp3** −3.29 Log2FC p-Val=1.34E-06), (**\*Ifi203** −2.38 Log2FC p-Val<1.00E-300), (**\*Gbp2** −2.99 Log2FC p-Val < 1.00E-300), (**\*Gbp2b** −3.25 Log2FC p-Val=3.00E-306),(**\*Gbp7** −1.77 Log2FCp-Val=4.27E-44),(**\*Gbp5** −2.75 Log2FC p-Val=2.37E-30),(**\*Gbp3** −1.97 Log2FC p-Val =7.54E-26),(**\*Ifi44** −3.42 Log2FC p-Val<1.00E-300), (**\*Ifi204**-2.50 Log2FC p-Val=1.15E-148),(**\*Il1b** −4.49 Log2FC p-Val<1.E-300), (**\*Casp4** −1.66 Log2FC p-Val=1.10E-23),(**\*Casp1** −0.85 Log2FC p-Val=2.57E-11),(**\*Irf7**-4.82 Log2FC p-Val=4.24E-72),(**\*Irgm1** −2.57 Log2FC p-Val=6.91E-276),(**\*Tank** −1.97 Log2FC p-Val= 5.77E-03), (**\*Il6** −3.91 Log2FC p-Val=2.84E-64),(**\*Tnf** −2.11 Log2FC p-Val=7.67E-59), (**Oasl2** −5.95 Log2FC p-Val<1E-300), (**\*Oas3** −2.51 Log2FCp-Val<1.00E-300),(**\*Oasl1**,-3.79 Log2FC p-Val=1.25E-85), (**\*Oas1a** −2.41 Log2FC p-Val =1.77E-62), (**\*Oas1g** −3.39 Log2FC p-Val =9.02E-32),(**\*Irf9** −2.22 Log2FC p-Val=1.70E-19),(**\*Myd88** −0.85 Log2FC p-VAL=1.97E-21), (**Card9** −1.4 Log2FC p-Val=3.68E-02), **Map1lc3a** (−2.1 Log2FC p-Val=4.10E-03), (**Trip6,** −1.3 Log2FC p-Val=8.38E-03).

We discuss the category of reverse tolerance separately, where in **Figure 10**, a hexagonal canvas plot shows the predominant pathways associated with this response. In addition, reverse tolerance involves acute repressed transcripts that acquire baseline levels in chronic conditions, most of which appear to involve regulated homeostasis in cell cycle/mitotic processes, which return to baseline levels during chronic inflammation. All these data will be further discussed, as the figures and results sections remain somewhat broad, and the data require a comprehensive interpretation of the totality of the dataset.

**Figure 10.**
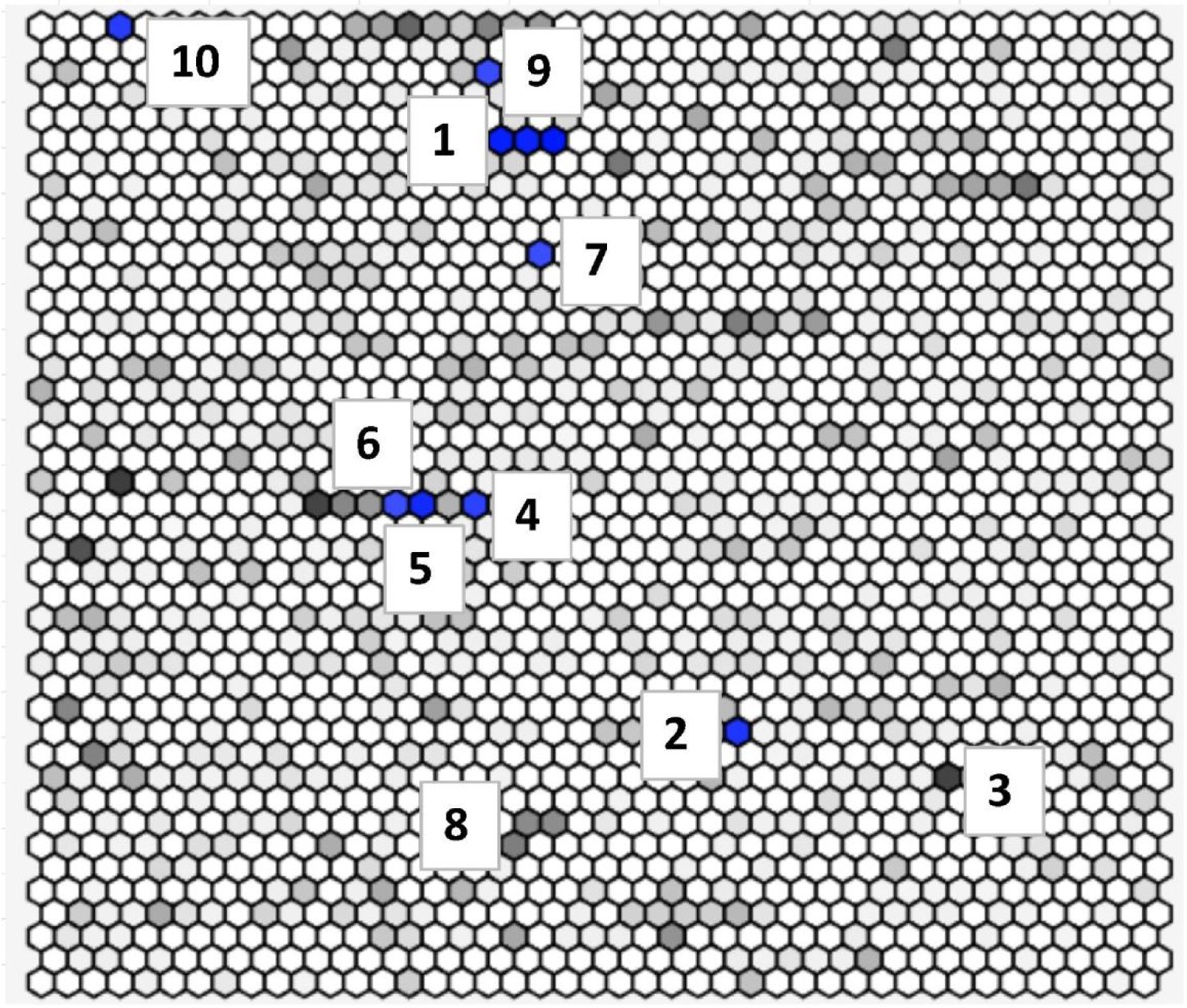
Chronic Inflammation and Reverse tolerance. A select number of DEGs were repressed in acute and restored to baseline. These are presented in a Hexagonal canvas plot generated from terms via the Reactome library. Each hexagon in the plot represents a single term. The brighter the color is, the greater the Jaccard similarity between the term and input gene sets. Similar gene sets are generally grouped together. The terms highlighted in blue most significantly overlap with the input query gene set. The numbers are provided for the spots corresponding to the tabulated list below.

## Discussion

This study clarified the phenomenon known as immune tolerance triggered by chronic inflammation or sustained LPS/TLR4 activation. It is imperative to align this response with a substantial portion of the human immune system, which ensures self-tolerance and the rapid attenuation of physiological immune responses postinfection to minimize pathological damage. This portion of the immune system (negative feedback) is an enormous force of magnitude. It is also likely responsible for limiting the efficacy of MAMP-based immune therapies and relapse from immune checkpoints (inhibitors ICIs). In cancer, this negative feedback substantiates a large part of the pathology, where tumors are surrounded by an overwhelming and aggressive negative feedback element that prevents the routine surveillance of self-malignant cells. Many terms are used to describe this process, such as “immune tolerance,” “immune exhaustion,” “loss of immunosurveillance,” “negative feedback systems,” “checkpoints,” and “chronic inflammation”. In this study, we used an in-vitro model to explore the effects of long-term sustained activation of the Toll-like 4 receptors (TLR4) (“chronic inflammation”) in macrophages. Macrophages play a formidable role in tumor immune escape (34), and as this work will show, chronic inflammation triggers an “M2” tumor-promoting phenotype, which enables the capacity of expansive, diverse myeloid-derived suppressor cells (MDSCs), dysfunctional CD+8 T cytotoxic cells (CTLSs), dysfunctional T suppressor cells (Tregs) and dysfunctional NK cells within the tumor microenvironment (TME) (35, 36).

In this study, chronic inflammation led to bidirectional DEGs that fell into eight classical patterns, many of which were not subject to either tolerance or reverse tolerance. (See Supplementary Table 1). The primary phenotype patterns associated with chronic inflammation include severe sustained loss of antigen recognition/presentation of MHC1, type I interferon signaling, antiviral, antibacterial host defense, IL-1αβ/TNFα signaling, acute phase proinflammatory cytokines, and complex metabolic disturbances. In contrast, chronic inflammation leads to the sustained overexpression of not only checkpoints (TIM3 and PDL1), but also suppressor of cytokine signaling 3 (SOCS3), IL-10, CCLs (2, 7, 12 +), extracellular matrix (ECM) degradative proteases and a host of other phenotypic changes as discussed below.

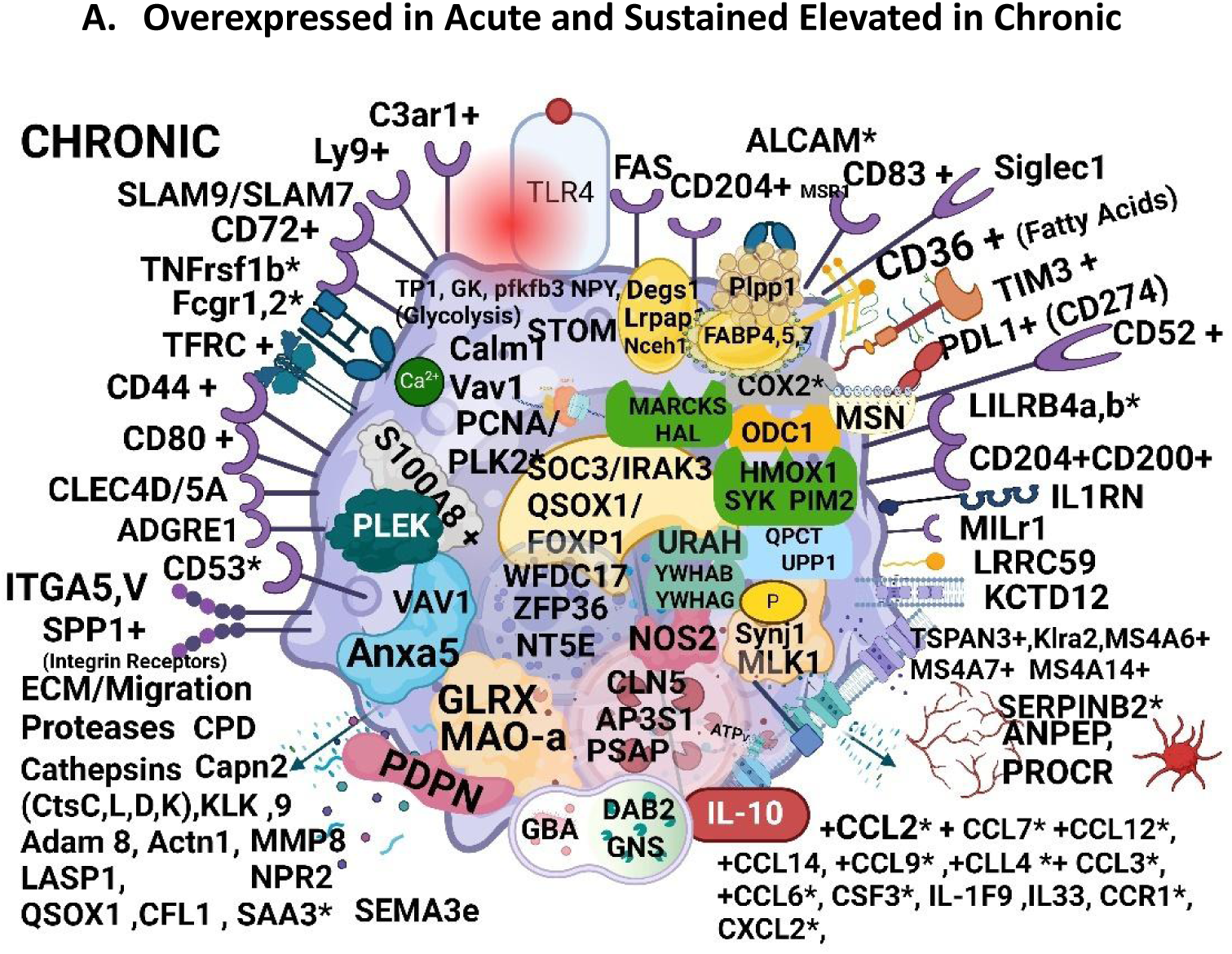

### Categories: Surface membrane glycoproteins, integrins and receptors

In accordance with therapeutic immune checkpoint targets, our data strongly align with acute and chronic inflammation, revealing the overexpression of the well-known PLDL1 and TIM3 checkpoint molecules, which truly take a back seat when the total number of negative feedback events reported as a result is considered. Throughout the discussion, the [*] denotes DEGs subject to tolerance, being elevated in acute and either completely reduced or slightly reduced but remaining above resting baseline levels, where we provide values in resting vs. 11-day chronic inflammation, which was similar to 7-day chronic inflammation. For example, we observed the overexpression of *CD274 (PD-L1) [+ 2.1 Log2FC, p-Val1.68E-03) along with its scaffold moesin (MSN) [+2.2 Log2FC, p-Val1.44E-02] a subunit of the ezrin, radixin, and moesin (ERM) complex (37, 38) and HAVCR2 (TIM-3), [+2.5 Log2FC p-Val, 1.24E-03] in chronic inflammation. These two checkpoints have been the subject of intense research, which, if blocked, are reported to correct dysfunctional CD8+ T cells and attenuate Tregs in cancer (39) for specific types of cancers (40). In the cell surface category chronic inflammatory doubling of osteopontin/secreted phosphoprotein 1 (SPP1) occurs at 7 and 11 days. SPP1 drives the modeling of the expression of surface chemotactic integrin receptors (α 4,9 β1 and α 9β4) which are highly characteristic of M2 TAMs and chronic inflammatory diseases and are expressed with PDL-1 and CTL4, with the capacity to usher in dysfunctional T cells, NK cells and those resistant to tumor immunotherapies tested against aggressive cancers (41–52). Like other checkpoints, therapeutically targeted blockade of SPP1 with monoclonal antibodies (*e.g.,* hu1A12, OPN-BsAb AOM1) reportedly ameliorates metastatic proclivity in human tumors (53, 54). It is useful as a risk biomarker (55, 56) and has been reported to spike in the initial stages of TLR4 tolerance in macrophages (57). Next, we found that chronic inflammation was associated with the overexpression of an integrin-related gene (ITGA5) and macrophage scavenger receptor 1 (MSR1), also known as CD204. CD204 encodes a fibronectin receptor involved in cellular adhesion and migration, and is overexpressed in invasive tumors contributing to chemotherapy resistance. It is useful as a prognostic indicator that is indicative of M2 TAMs and useful for guided monoclonal antibody-targeted treatments (58–62),; it is a membrane glycoprotein that is upregulated in acute/sustained chronic inflammation and is a feature of M2 −polarized TAMs. See review: (63)

In addition, the overexpression of Marcks (+3.0 Log2FC p-Val 7.88E-13) serves to guide integrin-dependent signal transduction pathways in macrophages (64) which are overexpressed in M2-TAM infiltrates in cancer patients with a poor prognosis (65). Many of the data are consistent with overexpressed immune checkpoints; these include the human leukocyte immunoglobulin (Ig)-like receptors (*LILRs) LIlrb4a (+2.1 Log2FCp-Val 6.50E-03)/Lilrb4b (+2.5 Log2FC p-Val8.02E-03) andS100A8+ (+7.3 Log2FC p-Val 4.31E-15) which together are reported to be abundant in M2 tumor-associated macrophages (TAMs), (66) driving tumor suppressor MDSCs, dysfunctional NK cells, T cells, and B cells (67–70). The LILRB receptor family also confers tolerance at the maternal–fetal interface, is involved in blocking NK cells, and establishes immunological memory to fetal antigens in a similar fashion to that of tumor-specific antigens (70, 71). The overexpression of LILRB most often coincides with extraordinarily high expression of the anti-inflammatory cytokines IL10, (+ 7.59 Log2FC p-Val 2.88E-14), CD83+ (72), and Tnfrsf1b (TNFR2 +) (3.0 Log2FC p-Val6.05E-14)(73) in inflammatory pathologies and in non-responders to ICIs for cancer treatment, such as lenvatinib plus anti-PD1 antibodies (74). These sustained changes in expression are also reflected by a rapid takeover of negative feedback subpopulations including tumor-infiltrating CD4(+) Foxp3(+) regulatory T cells (Tregs) marked with checkpoints (e.g. CTLA-4 and TIGIT), dysfunctional CD8(+) T cells, and a host with an inability to recognize and destroy self-malignant cells (75–78).

Some of the markers of chronic inflammation, which are provoked during acute inflammation and attenuated during chronic inflammation, remained **slightly above the resting baseline**, suggesting that they could still play a role in chronic inflammation, so we will discuss these markers. Some of these include the sustained overexpression of *C3ar1 (+ 1.4 Log2FC p-Val 1.90E-02a) a marker of self-tolerance (79), which is associated with dysfunctional NK cells, dendritic cells, T and B cells, expressed TIM3 and PD-1 in advanced tumors (80–85), chronic inflammation and sepsis (86, 87), and are deemed as effective antitumor ICI targets (88, 89). Another example is the activated leukocyte cell adhesion molecule (*Alcam); resting vs. chronic (1.3 Log2FC p-Val 9.52E-02) which docks to CD6 on T cells, B cells, NK cells, immune cells, fibroblasts, and endothelial cells conferring dynamic tumor immune escape and metastatic risk (90). Even though these effects appear very mild in relative magnitude within this study, they are being explored in tumor immune therapies such as the antibody‒drug conjugates praluzatamab ravtansine or CX-2009 (Alcam), which are undergoing phase II clinical trials (91). Others in this category, including macrophage scavenger receptor 1 [CD204(+) *] (+2.0 Log2FCp-Val 4.0E-03) and CD200(+)) (+3.1 Log2FC p-Val 3.11E-04), which are highly expressed in M2 TAMs (92, 93),; associated with chemo resistant cancers (94) and infiltration of dysfunctional CD8+ T cells (95–98) with increased *CD53+ (99), are also involved in tumor immune escape and susceptibility to recurrent viral, bacterial, and fungal infections (100).

Larger changes evoked in acute inflammation (subject to minor tolerance) but significantly elevated above baseline include the interleukin 1 receptor antagonist [*+Il1rn] (3.9 Log2FC p-Val 3.04E-04), a transcript subject to genetic mutations involved in cancer (101, 102), inflammatory pathologies in particular asthma (103, 104), and viral infections such as SARS-CoV-2 (105). We also observed chronic overexpression of CD73(+), also known as 5’-Nucleotidase Ecto (NT5E) (+3.5 Log2FCp-Va l 4.73E-05), a proposed therapeutic target to reestablish antitumor CD8(+) T-cell NK cells, increase the efficacy of ICIs (106), and restore the loss of T-cell receptor signaling and interferon (IFN) (Types I & II) pathways (107). Chronic inflammation triggered the overexpression of CD44+(108, 109) (2.2Log2FCp-Val9.29E-03), SLAMF7(+) (CRACC; CS1; CD319), and +4.6 Log2FC p-Val 1.30E-13), the latter of which is a well-known negative feedback restraint that placates septic shock (110) and reduces *TNF-alpha which was also found in this study (−2.11 Log2FC p-Val=7.67E-59) in LPS vs chronic (111). Therapeutic anti-SLAMF7 monoclonal antibodies such as elotuzumab have been used to treat multiple myeloma by reestablishing antibody-dependent cellular cytotoxicity (ADCC) and NK cell-targeted ligation (112). We also observed the overexpression of SLAMF9 (+) (CD84-H1; SF2001; CD2F10) to a lesser degree (+2.2 Log2FC p-Val2.89E-04) where both SLAM-family receptors are now considered viable future targets for cancer immunotherapy (113). Macrophage scavenger receptor 1 (MSR1), also known as CD204, is a membrane glycoprotein that is upregulated during acute/sustained periods and is a feature of M2 −polarized TAMs; see review: (63)

### Category signaling: SOC3

The data revealed that all of these changes are tantamount to heightened SOCS3 signaling, which inhibits JAKS 1,2 TYK2 and then binds to a gp130 receptor to invoke negative regulation of *IL-6 (chronic vs. LPS) (−3.91 Log2FC p-Val 2.83989E-64), G-CSF (Csf3) (−3.71 Log2FC p-Val 3.2812E-117), andCD40(114), whereas the opposite occurs with the increase in IL-10 (+7.6 Log2FC p-Val 2.88E-14) (115–117) and SOCS3 (+4.9 Log2FC p-Val 8.73E-20). These changes occur with a step-up in the expression of serine/threonine-protein kinase (Pim-2) (+ 3.0 Log2FC p-Val 2.31E-06), which is now subject to investigational drug discovery for the treatment of hematologic cancers, including acute myeloid leukemia, mantle cell lymphoma, and anaplastic largecell lymphoma (ALCL), showing greater efficacy when combined with ICIs (118–120).

### Category-Response ECM/ Cathepsins

Chronic inflammation triggers very high levels of S100a8 (+7.3 Log2FC p-Val 4.31E-15), a calcium-binding protein that promotes neutrophil migration in chronic virus-induced inflammatory pathologies (e.g., COVID-19), (121) septic shock and cancer and is often expressed with PD-L1 (122–125). These immunosuppressive forces promote ECM matrix degradation, aiding in cell motility (126) which in this work also involves the upregulation of cathepsins C, D, and L, respectively (Ctsc +2.3 Log2FC p-Val8.79E-06), (Ctsd +2.6 Log2FC p-Val 3.17E-03), (Ctsl +3.1 Log2FC p-Val2.31E-08), as well as Adam 8 (+ 2.3 Log2FC p-Val1.27E-04), carboxypeptidase D (CPD), (+3.9 Log2FC p-Val 8.99E-06) and kallikrein-related peptidases (Klk9) (+7.2 Log2FC p-Val 7.03E-10).The upregulation of cathepsins in this study, coincided with the underexpression of their inhibitors (cystatin c (CST3)) (−2.5 Log2FC p-Val 4.88E-05) where opposing synergistic changes are described in rapid tumor growth, immune escape, and pathological chronic inflammation (127–130). Overexpressed cathepsins, which are typically indicative of an aggressive malignancy, are used as risk biomarkers (130, 131) and serve as targets for anti-CTS tumor therapies that reinvigorate NK function, revert the antitumor M1 phenotype, attenuate MDSCs, enable an antitumor CD8+ T-cell response (132–137) and increase the efficacy of antiviral therapies (e.g. SARS-CoV-2) (138). The co-overexpression of cathepsins, along with ADAM8 and CPD, is a formidable force in driving pathological roles in chronic inflammation, asthma, heart disease and cancer (126, 139–143), which is highly aligned with the breakdown of the ECM with mobility chemokines,particularly CCL2(+6.0 Log2FCp-Val 8.61E-19) (126). Similarly, the data in this study show that chronic inflammation is associated with sustained elevation in a series of CCLs, most significantly CCL2 and CCL7 (+ 5.8 Log2FC p-Val 1.79E-06), which are both drivers of TAM M2 trafficking into tumors in synergy with MDSCs (144). These combined forces are largely responsible for resistance to tumor immunotherapies (145) and poor clinical outcomes (e.g. colon, uterus and lung) (146–148). The increase in CCL2 also paralleled that of CCL12, (+5.8 Log2FC p-Val 1.36E-03) and CXCL2 (+4.04 Log2FC p-Val 1.24E-14) which amplify the CCL2/CCL12-CCR2 axis; one of the most pivotal trafficking patterns associated with the infiltration of MDSCs tagged with CCR2s (e.g. basophils, DCs, NK cells and T cells) in cancer (36). This axis is a well-established avenue for antitumor therapies and justifies the development of CCL2/CCL12 monoclonal antibodies that increase the efficacy of anticancer vaccines (149). We also observed the overexpression of QPCT (+4.7 Log2FC p-Val1.98E-10), which is involved in the recruitment of CCL2/CCR2 signaling, under chronic conditions (150).

### Category-Metabolism

The data in this work highlight the major metabolic disturbance in chronic inflammation reflected by the overexpression of CD36(+) (leukocyte differentiation antigen) (+2.8 Log2FC p-Val 1.64E-02) known as fatty acid translocase (FAT) which aids in the transport of fatty acids and oxidized lipoproteins into the cell, as well as a known response to the activation of Toll-like receptors (e.g. TLR4,6,7) (151), all of which are found in M2 TAMs (152). The CD36(+) marker on immune cells in the TME is reflective of greater risk for metastasis, inferior chemotherapy response, and the appearance of abundant Tregs (153, 154) which is characteristic of dysfunction in lipid metabolism (155) and is often listed as a “hub gene” in data analytics that aims to correlate biomarkers with poor predictive prognosis and tumor immune escape (156–158).

The presence of CD36(+) also aligns with parallel elevation in the fatty acid binding proteins Fabp7 (+3.6 Log2FC p-Val3.40E-07), Fabp5 (+2.7 Log2FC p-Val 1.73E-05), Fabp4 (+ 2.4 Log2FC p-Val 1.57E-02) and the reduction of lipoprotein lipase Lpl (−2.5 Log2FCp-Val 1.40E-10). These effects aid in the accumulation of triglycerides often reported in immune cells to aid in escape and are instrumental in the pathology of hypertriglyceridemia, hypercholesterolemia and atherosclerosis, which are reversible by LPL agonists, such as pioglitazone NO-1886 (159, 160). LPL agonists with anti-CD36(+) therapies are also used to augment the efficacy of anti-PD-1 therapies (161). We also observed a sustained reduction in plasma phospholipid transfer protein (PLTP), a member of the lipid transfer/lipopolysaccharide-binding protein (LT/LBP) family involved in lipoprotein metabolism, a deficiency reflective of the anti-inflammatory CD4+ Th2 phenotype (162), and cholesterol accumulation in disease-related macrophages (163). Overall, increased fatty acid uptake into macrophages (CD36+) could also generate substrates for *COX2 Ptgs2 (+3.8 Log2FC p-Val 2.73E-04) (152). All of these systems (as systems that are overexpressed or highly active) have been the subject of extensive research in the field of immune oncology. See Reviews: (151, 164, 165). In terms of fat metabolism, chronic inflammation is associated with minor acute and chronic sustained elevations in spleen tyrosine kinase (SKY) which is essential for TLR4 activation by host-derived DAMPs and plays a role in fatty acid deposition in atherosclerosis (166, 167). The use of SKY inhibitors, such as R788 (fostamatinib), in conjunction with anti-PDL1 provides a synergistic effect leading to complete tumor regression and durable antitumor immunity in mice bearing small tumors (168).

Metabolic changes in the glycolysis/pentose phosphate pathway were also noted where chronic inflammation triggered undulations of Aldoc (−1.2 Log2FCp-Val 1.17E-03), and Aldoa (−1.8 Log2FC p-Val 1.70E-02) which are needed to convert fructose 1,6-bisphosphate, into dihydroxyacetone phosphate (DHAP) and glyceraldehyde 3-phosphate (G3P), which are needed for glycolysis. We also observed major collective losses in mitochondrial OXPHOS genes, such as mt-Nd4 (−2.4 Log2FC p-Val 1.41E-04), mt-Nd6 (−2.3 Log2FC p-Val 2.82E-03), mt-Te (−2.5 Log2FC p-Val 1.79E-03), mt-Cytb (−2.2 Log2FC p-Val 2.54E-04)ATp5e, ATp5g2, and the OXPHOS III-IV super complex assembly factor Higd2a (−1.9 Log2FCp-Val 4.43E-03) (169, 170). Our work correlates well with research reported in the literature, where one of the major changes in this work is the loss of NDUFS7 NADH:ubiquinone oxidoreductase core subunit S7, an essential element of complex I which is often observed in the M2 suppressive phenotype (171). These changes are compounded by major losses of Atpif1, which could render the collapse of the structural stability of electron transport (172) worsened by a loss of CARD19 (−2.3 Log2FC p-Val 4.04E-03) housed within mitochondrial cristae (173). We also observed a severe tolerance-related nonresponse in *Cmpk2 (−3.97 Log2FC p-VAL 1.79E-280), which is believed to arise from faulty IRF1 signaling, and is also lost to tolerance, which is required for mtDNA synthesis and the NLRP3 inflammasome complex (174, 175). Mitochondrial defects associated with acute inflammation have been reported consistently throughout the literature (57) where dysfunctional OXPHOS is also associated with an increase in PLK2 (also found in this study), which needs to be further investigated to determine how these two events are linked (176–178).

One of the most unexpectedly overexpressed DEGs in chronic inflammation was NOS2 (NOS2 +5.5 Log2FC; p-Val =5.7 ×10-34), which was associated with the opposite (severe loss) of SOD2 (−2.51 Log2FC p-Val= 2.15E-164) in chronic inflammation, promoting the rapid formation of ONOO-toxic radicals. This was highly unexpected because the M2 phenotype is believed to be devoid of NOS2. NOS2 is highly sustained in chronic inflammation, as evidenced by a gain in secondary compensatory detoxification systems that neutralize NO and ONOO-radicals such as the redox system S-nitrosylation defense system iron-dinitrosyl thiolates (MT1) (179, 180), S100A8, (181) QSOX1, (182, 183) glutaredoxin (Glrx), and heme oxygenase (Hmox1) (184–186). Interestingly, most of these response genes, with the exception of NO2, are reportedly overexpressed at tumor immune checkpoints and demarcate aggressive drug-resistant cancers (187, 188), which are occupied by dysfunctional CD8+ T cells (189, 190).

Changes in metabolism have also been observed in the regulation of iron metabolism. These include a sustained rise in the transferrin1 receptor which is the gateway for iron transit in and out of cells, occurring with simultaneous loss of the intracellular chelator Hscb (HscB iron‒sulfur cluster cochaperone), Iscu iron‒sulfur cluster assembly enzyme, Cisd3 CDGSH iron sulfur domain 3 and Fth1 ferritin heavy polypeptide 1; which is consistent with reports suggesting that faulty iron storage capacity is a hallmark of M2 macrophages with these traits (191). The loss of Fth1, an iron storage protein, observed in this study suggests a greater propensity for metal-mediated oxidative toxicity concomitant with losses in FXYD5 and 2 (ion-transport regulators), events that may drive immune-escape and Treg infiltration in colon cancer, which aligns with the work of Zhang *et al*., 2022 (191).

#### -Signature Checkpoint Genes

One of the most marked effects of chronic inflammation in this study was sustained elevation of a series of small proline-rich proteins (SPRRs): SPRRs 2B, 2E, 2D, 2F, 2G, 2H, 2I, 2J, and 2K. These were the most dramatic shifts in the transcriptome. Very little research has been conducted on the functional aspects of these proteins. They are also expressed in the dermal cutaneous barrier, and are stimulated by TLR4/MYD88 agonists, they are known to have potent bactericidal properties against methicillin-resistant *Staphylococcus aureus* (MRSA) (192), are linked to chronic inflammatory diseases (193) (194), and respond to noxious stimuli such as cigarette smoke (21). We also observed a chronic inflammation-mediated rise in several transcripts to which little is known, such as histamine ammonia lyase (HAL) and STFA2 and 3 (Stefins). Sprr2b (+7.47 Log2FC p-Val 1.34E-07), Sprr2c-ps (+7.13 Log2FC p-Val 1.13E-08), Sprr2d (+7.56 Log2FC p-Val 1.28E-10), Sprr2e (+7.44 Log2FC p-Val2.05E-11), Sprr2f (+7.56 Log2FC p-Val 1.12E-13), Sprr2g (+ 8.42 Log2FC p-Val 2.98E-06), Sprr2g (+ 7.14 Log2FCp-Val 3.28E-15), Sprr2h (+8.12 Log2FC p-Val 5.11E-13) Sprr2i (+7.56 Log2FC p-Val 8.30E-12), Sprr2j-ps (+8.10 Log2FC p-Val1.13E-11), Sprr2k (+7.10 Log2FC p-Val 2.28E-05), Stfa3 (+7.00 Log2FC p-Val 1.89E-14), Hal (+4.32 Log2FC p-Val 1.04E-14).

In the same manner, other overexpressed DEGs at 7 and 11 day chronic inflammation would include: 1) tetraspanin (e.g Tspan 3), a facilitator of microbial binding, overexpressed in human cancers(195, 196), an enabler of migration/adhesion, (197) tumor immune escape (198) chronic viral infections (199) and inflammatory pathologies (200–205); 2) Qsox1 quiescin Q6 sulfhydryl oxidase 1 (QSOX1); a driver of matrix degradation, tumor invasion, the latter subject to monoclonal antibody inhibitors in models of breast cancer (206–208); 3) prodoplanin (PDPN) a driver of epithelial-mesenchymal transition (209), resistance to trastuzumab, dysfunction in NK (210), CD8(+) T cells, presence of M2 CD204(+) TAMs/CAFS, reduction in interferon-gamma, granzyme B and perforin-1 (211) and found in progressive cancers (212–214); 4) Plaur (CD87 receptor for urokinase plasminogen activator) (215, 216) and other genes associated with the upregulation in acute/sustained during chronic inflammation, a feature of the M2 polarized TAM: See review: (63). For most systems in this study which align with general reports on the observation of M2 TAMs, TANS, immune infiltration of immunosuppressive MDSC-derived dysfunctional NK and CD8(+) T cells, which are associated with a greater risk of viral/bacterial infection, higher pathogen titers, and a lack of response to anti-PD-L1 therapy and overall worse survival in cancers including Adgre1 (217) QPCT1 and Ms4a7 (218, 219) Fcgr1 Fc receptor, IgG, high affinity (CD64(+)) (220), Msr1(221), CALM1 (222), TPI1 (223), SRGN, (224, 225) Anxa5(226) (227, 228) (229–232), Wfdc17 (233), and IL1F9 (234–236).

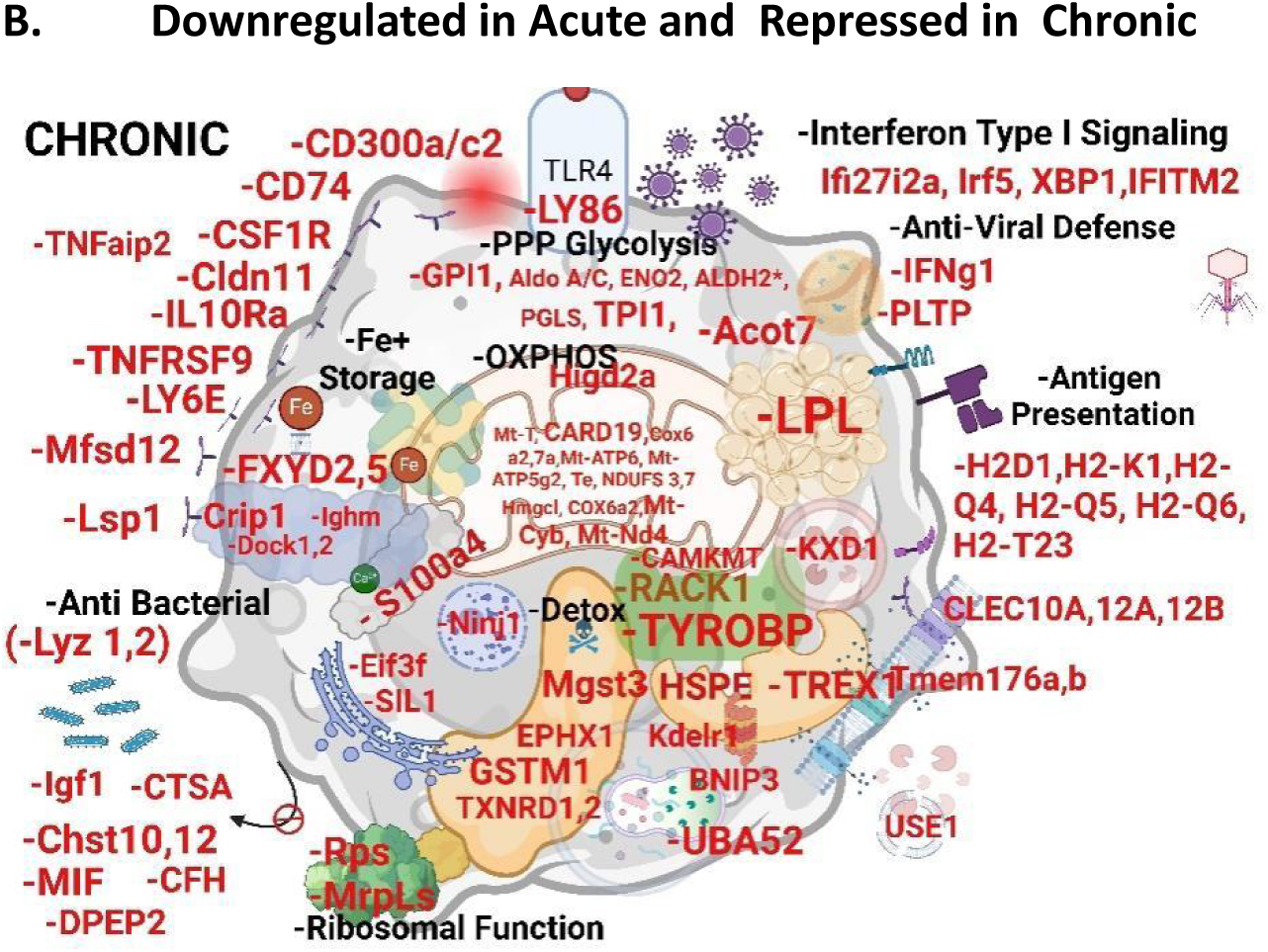

Category -Antigen Presentation: The most striking loss involved in chronic inflammation in this study was a sustained collapse of histocompatibility genes [H2-Q4, H2-Q6, H2-T23, H2-D1, H2-T22, H2-Q4, H2-T23, H2-T24, H2-T22, H2-K1, and H2-Q5] which play a role in antigen presentation in the host defense against self-virally infected or malignant cells and are also essential for vaccine efficacy to program acquired T cytotoxic cell and B-cell antibody responses. Collectively, these losses handicap the MHC class I ER pathway, which is otherwise needed to drive T-cell-mediated toxicity, a known ortholog of human HLA-A (class 1, A, B and C) to which a loss leads to greater susceptibility to infectious diseases (237). This concept was recently demonstrated by Xiong *et al*., who studied signaling systems involved in the impediment of IFN-gamma signaling (also reduced in this study), and reported that the loss of MHC-1 H2-K1 and H2-D1 (and phagocytosis) concomitant with the loss of CD8(+) T cytotoxic infiltration is required to attenuate tumor growth in mice (238). The data in our study revealed major losses in both MHC1 and type I IFN-related genes, severely reducing the ability of the host to present the antigens required to foster T-cell immunity, particularly against cancer and infection (239). The loss of H2-Q5 histocompatibility 2 (Q region locus 5 in humans), is also observed at the maternal–fetal interface where silencing of the innate/adaptive immune system is needed to prevent NK and T cells via CD94/NKG2 receptor-mediated toxicity (240). Moreover, CD40 agonists can increase HLA-DR (+) levels, propagate the transition of macrophages to the M phenotype, suppress human pancreatic cancer (241), and are a positive prognostic indicator in patients with cancer (242). In contrast, the loss of HLA-DR and TNF-α is observed in sepsis patients with an impaired immune response (243).

### Category-Interferon Signaling

The data show that chronic inflammation associated with tolerance is highly linked to enormous losses in a large number of interferon-related genes including IFIS (27, 30, 44, 44I, 25, 203, 203-ps, 202b, 204, 205, 207, 209, 211, 213, 2712a) IFITs (1, 2, 3, 3b, m2) IRF2 bp2 IFRD1, IFNGR1, IFIH1, IRFs (5, 7) IGTP, and ISG20. The loss of IFN-stimulated genes (ISGs) prevents host defenses against viral/bacterial infections and cancer (244), which aligns with the therapeutic use of interferons for decades to treat cancer (245). One of the most remarkable losses we see here is that for the class of IFITMs such as *IFIT1 (−6.31 Log2 FC p-Val <1.00E 300), *IFIT2 (−2.64 Log2FC p-Val= 3.09E-101) *IFIT3 (−4.97 Log2 FC p-Val <1.00E-300), *IFIT3b (−5.08 Log2FC p-Val=3.06E-164), *IFITM3 (−2.02 Log2FC p-Val=7.29E-27) *IFI204 (−2.50 Log2FCp-Val=1.15E-148), *IFI211 (−2.12 Log2FC p-Val=5.84E-97),*IRGM1 (−2.57 Log2FC p-Val=6.91E-276), *IFI44l (−2.99 Log2FC p-Val=2.25E-121),*IFIT1bl2 (−4.00 Log2FC p-Val=4.78E-31), etc., which as host effectors are required to launch defense against gram-negative pathogens, (246), preventing viral replication (247) and virulence (248), and allowing greater susceptibility to Hep C (249), Ebola, influenza, SARs-COV and West Nile (250). The data on chronic inflammation (tolerance) establish a sustained concomitant reduction in type I IFN receptor transporter protein 4 (251) Rtp4 (−5.97 Log2FC −Val<1.00E-300) and the interferon-inducible Ly6e lymphocyte antigen 6 complex, locus E, which is repressed in acute and chronic Ly6e (−2.8Log2FC p-Val=1.83E-05), and is highly involved in antiviral immunity involving B-cells, T cells, dendritic cells, and macrophages for defense against HIV-1 (252) SARS-CoV-2 (253) and Middle East respiratory syndrome coronavirus (MERS-CoV) (254–256). These losses appeared to be very prominent and occurred in tandem with major losses across the board for a host of IFN inducible genes, including XBP1, a deficiency of which leads to a greater bacterial burden of *Francisella tularensis* in animals which is essential for TLR to IFN-g signaling via the IRE1–XBP1 pathway (257, 258). In general, the loss of many IFN systems in macrophages results in an M2 phenotype, (259) and impairs host defense against a wide variety of pathogens including DNA viruses, bacteria, *Staphylococcus aureus, and M. bovis* infections (260–264) in addition to *Mycobacterium tuberculosis* (265) and HIV (266). The data also revealed chronic inflammation-mediated tolerance induced losses in *Isg15 (−5.56 Log2FC p-VAL>1.E-300), *Isg20 (−3.71 Log2FC p-Val=1.91E-31) and (p62), and *Sqstm1 (−2.41 Log2FC, p-val=1.84E-220) constitute a major arm of the antiviral type I interferon-conjugated autophagy clearance response (267–271). P62 disruption is associated with a lack of proper maintenance of protein degradation, involving not only cancer but also age-related neurodegenerative diseases (272). Therapeutic plasmids, such as elenagen, which encodes p62/SQSTM1, have been tested as anticancer DNA vaccines for treating aggressive solid tumors (273, 274). Our work corroborates the recent work of Makjaroen, J. *et al*., 2023 who demonstrated that a second dose (repeat) of LPS in macrophages was associated with a reduction in glycolysis, OXPHOS, ETC transcripts, antiviral responses, ISG15, and loss of antigen presentation pathways (275). We also observed a tolerance nonresponse for *Aoah (−3.83 Log2F p-Val=3.70E-58), which is a lipase acyloxyacyl hydrolase, with defects or silencing in mice resulting in greater susceptibility to *Pseudomonas aeruginosa* (276). LPS is deacylated/ detoxified by AOAH and in hepatic Kupffer cells when its loss is believed to increase the endotoxic potential of LPS because of its presence remaining in systemic circulation (17).

The data in this work also show chronic inflammation associated with a sustained concomitant reduction in a series of interferon-induced 2’-5’-oligoadenylate synthetase transcripts 1a,1 g,3 and 1; *Oasl2 (−5.95 Log2FC p-Val<1.00E-300), *Oas3 (−2.51 Log2FC p-Val<1.00E-300), *Oasl1(−3.79 Log2FC p-Val 1.25E-85), *Oas1a (−2.41 Log2FC p-Val 1.77E-62), *Oasl2 (−6.70 Log2FC p-Val 8.57E-45), *Oas2 (−7.61 Log2FC p-Val 5.03E-44), *Oas1g (−3.39 Log2FC p-Val 9.02E-32) *Oasl2 (−6.66 Log2FC p-Val 2.87E-26) well-known host defenders against viral RNA in part by blocking replication (277, 278) and the series of Schlafen genes (SLFNs), required for acquired immune response to microbial pathogens which is lost after acute phase response in macrophages by LPS (279). Our work also demonstrated that this was the case with an acute phase increase and no response to chronic inflammation *Slfn1 (−5.90 Log2FC p-Val=2.16E-53) *Slfn4 (−2.76Log2FC p-Val 3.87E-291) and *Slfn2 (−2.51 Log2FC p-VAL 4.42E-113). In theory, the sustained loss of SLFN2 during chronic inflammation impairs T-cell- and monocyte-mediated immunity (280), thereby substantiating greater susceptibility to bacterial and viral infections (281) and cancer (282).

The collective losses in the IFN response system during chronic inflammation also include deficiency in (TRAF family member-associated NF-kappa-B activator) TANK, along with *Nfkb2 (−2.01 Log2FC p-Val 6.37E-89) and *Irf7 (−4.82 Log2FC p-Val 4.24E-72). TANK is an adaptor protein that regulates Tank binding kinase 1 (TBK1), which phosphorylates IRF7 and translocates to the nucleus to promote the type I IFN response. Tolerance-mediated loss of IFN Type 1 signaling by sustained repression of IRF7 handicaps the antiviral regulator of the type I IFN host defense system (283). This IRF7 deficiency alone confers greater susceptibility to viruses and cancer, a dominant overtaking of MDSCs and M2 macrophages, and rapid tumor growth and metastasis (284–286). In human cancer patients, there is a direct association between high Irf7 expression and prolonged survival with a deficiency of a negative prognostic indicator (287, 288).

Similarly, tolerance in chronic inflammation is associated with the loss of 1) interferon-induced GTP-binding protein (Mx), which is involved in host resistance against RNA viruses (289, 290) 2) the IFN-inducible gene; bone marrow stromal cell antigen 2 (BST2), which is required for antiretroviral defense of the innate immune response (291–293) and 3) a reduction in Rsad2 (−4.13 Log2FC p-VAL >1×10--10)/viperin, another interferon-inducible protein involved in host defense against DNA and RNA viruses, oncogenic viruses such as simian virus 40 (SV40) large T antigen (TAg) (294) and measles (295). Tolerance mediated response losses also involved several IFISs, such as *IFI44 (−3.42 Log2FC p-Val <1.00E-300), *IFI203 (−2.38 Log2FC p-Val<1.00E-300), and *IFI208 (−3.65 Log2FC p-Val 6.77E-66). This coaligned with further loss of receptor transporter protein 4 (RTP4), which is involved in restricting viral replication including that of the human coronavirus SARS-CoV-2 (296), and loss of poly(C)-binding protein 2 (PCBP2) which is involved in antiviral immunity (297). The data also revealed that chronic inflammation is associated with a sustained reduction in the host defense transcript Sp140 nuclear transcript (GSK761) (298) (299) where Sp140(−/-) mice are susceptible to various bacterial infections (300) with higher SP140 expression favorable in head and neck squamous cell carcinoma, involving a greater response to ICIs, pathology involving M1 TAM phenotype transition and infiltration of functional CD8(+) T cells in various tumor models (301, 302). Depletion of SP140 in macrophages is associated with compromised microbe-induced activation and SNPs markedly suppress innate immune response genes (303).

#### -IRG1/ACOD1

The data revealed that chronic inflammation is associated with a sustained reduction in immune-responsive gene 1 (IRG1) (304) which is also reflective of a dysfunctional TCA cycle and OXPPHOS, as previously discussed. A tolerance-mediated response loss of “Acod1” aconitate decarboxylase 1, *Acod1 (−4.18 Log2FC p-VaL <1.00E-300) is well known to be involved in the pathology of inflammatory diseases such as atherosclerosis, infection (305–308), pulmonary fibrosis, herpes simplex virus-1 and −2 (HSV-1 and HSV-2), vaccinia virus and Zika virus, various bacterial infections (309) insulin resistance and lipidemia (310). There has been considerable interest in the development of itaconate derivatives, such as dimethyl itaconate (DI), 4-octyl itaconate (4-OI), and 4-ethyl itaconate (4-EI)(311), which can attenuate chronic inflammatory disorders, such as osteoarthritis, airway inflammation and prostatitis (312), and Behcet’s uveitis (313). The role of itaconate in cancer suggests that a high rate of ACOD1 product formation is associated with antitumor effects with the ability to augment the response to anti-programmed cell death protein immunotherapy (314).

### Pathogen–host response

#### -Lysozymes

The data also revealed that chronic inflammation is associated with a sustained reduction in lysozyme (Lyz1, Lyz2), resulting in a major loss of first-line defenders present in mucosal secretions involved in innate immunity against bacteria, viruses, and fungi. The use of exogenous lysozyme in medical applications against infectious diseases has been proposed, (315) because of its ability to destroy viruses and bacteria by disrupting pathogen replication through the electrostatic binding of nucleic acids and lytic properties (316). The data show that chronic inflammation is associated with a sustained reduction in serum amyloid A3 (M-SAA3), which is needed to provide natural innate destruction against host infections such as *Staphylococcus, Streptococcus*, and *E*. *coli* (317). Deletion of Saa3 increases susceptibility to infections such as *Pseudomonas aeruginosa* in mice (318) manifests as major defects in the innate immune response (319) and is associated with greater mortality from influenza (320).

### CD14/ CLEC4E

The data also revealed a loss of tolerance in response to CD14 (LPS-TLR4 coreceptor)(321), and a loss of response to PAMPS, DAMPS, or MAMPS, which bind to TLR4 receptors. These losses were associated with the C-type lectin receptor CLEC4E, *Clec4e (−3.64 Log2FC p-Val <1.00E-300), a transmembrane protein highly dense in APC myeloid cell lineages, including dendritic cells, which are required for pathogen recognition (322) DAMPs/MAMPS (322), and LPS, which binds to its CRD domain (322, 323). A deficit or SNP mutation in CLEC4 confers greater susceptibility to a diverse range of infectious diseases, including *M. tuberculosis, Streptococcus pneumoniae,* sepsis, meningitis*, Tannerella forsythia, Corynebacterium*, a variety of fungi and parasites (322, 324, 325). Loss of both TLR2-/ and CLEC4 (also found in this study) can amplify the loss of antigen-presenting proteins and increase susceptibility to *C. albicans* infection (326). By mechanism of action, CLEC4s are essential for acquired immunity, a deficiency linked to inadequate affecter B cells, and T lymphocyte function (327), and their expression at high concentrations is an attribute of antitumor M1 macrophages (328); see Review (323).

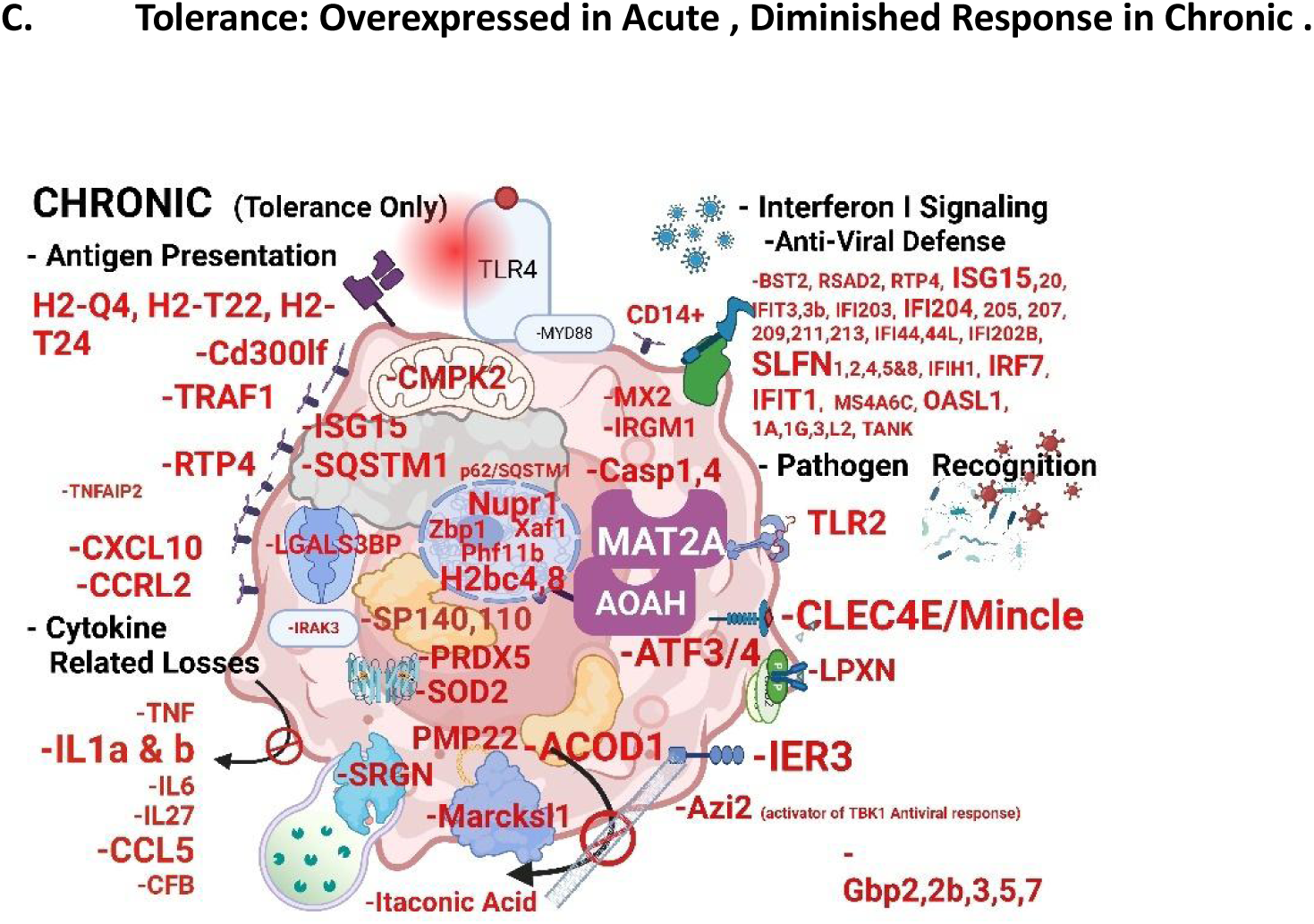

### G-CSF (Csf3)

The data also revealed that chronic inflammation is associated with a tolerance-mediated reduction in Csf3, colony-stimulating factor 3 (granulocyte) (G-CSF) *Csf3 (−3.71 Log2FC p-VaL 3.28E-117), which still remained 10 times greater or (+3.4 Log2FC p-Val 4.55E-02) than resting during chronic inflammation. Csf3 is instrumental in the mobilization of neutrophils, the loss of which can be overcome by the use of recombinant peptides commonly used to treat congenital and acquired neutropenia, chemotherapy-induced neutropenia, and fungal infections (329). Paradoxically, with respect to cancer itself, G-CSF-producing malignant cancers are associated with poor clinical prognosis (330) in high-grade tumors (330, 331) and have the capacity to generate immunosuppressive neutrophils (TANs) (35); however, recombinant G-CSF is used in cancer patients to reverse chemotherapy/radiation-induced neutropenia and has been proven safe and effective (332–334). This could introduce a double-edged sword in the treatment of cancer, as evidenced by G-CSF causing accelerated tumor growth, greater tumor vasculature, and increased immune suppressor and endothelial cell-differentiating cells in tumor-bearing mice (335). The data in this study also revealed tolerance-mediated chronic inflammatory full loss of complement factor *Cfb (−19.02 Log2FC p-VAL 2.87E-25). In terms of loss, it has been associated with poor clinical outcomes and lower survival rates in patients with lung cancer (336), most of which are associated with a greater risk of age-related macular degeneration (336, 337).

### Tumor Necrosis Factor Signaling / Interleukin 1 αβ

The data in this work show that chronic inflammation reduced in tolerance involves an ablated response to LPS through many proinflammatory responses including*Traf1 [4.15 Log2FC p-VAL 7.95E-197], TNF-alpha *Tnf (−2.11 Log2FC p-Val 7.67E-59), TNFAIP2,and TRAF2, which encode genes for the TNF receptor (TNFR)-associated factor (TRAF) protein family and are lost to Tnfrsf1b, Tnfrsf22, Tnfrsf23, and Tnfrsf26, however, the last four transcripts were sustained over the resting baseline control. The slight increase in Tnfrsf1b (TNFR2) above baseline during chronic inflammation could be associated with the infiltration of CCR8+ MDSCs, which, if blocked, can augment the efficacy of anti-PD1 therapies (338–340). The loss of TNF-alpha systems in this study is profound, which would render a greater risk for infection and tumor-promoting immune phenotypes: M2 TAMS and N2 TANs concurrent with dysfunction of antitumor activated CD4(+) and CD8(+) lymphocytes (341). We also observed tolerance to significant losses in IL-1/TLR signaling. This leads to a poor response to ICIs, and the acceleration of chemical and oncogene-induced cancers (342). In this study, we observed almost complete losses in IL-1s, including *Il1a (−5.16 Log2FC p-VAL <1.00E-300), *Il1b (−4.49 Log2FC p-VAL < 1.00E-300), and interleukin-1 receptor-associated kinase-3* Irak3 (−18.27 Log2FC p-Val 1.87E-24), with sustained elevation in Il1rn (Il1 receptor antagonist) and changes congruent with the overall loss of IL function (343) Irak3 observed in chronic inflammatory conditions such as rheumatoid arthritis(344).

### Leupaxin

LPS tolerance also involves chronic loss of leupaxin (LPXN), a protein not subject to a great deal of research but has been reported to play a functional role in humoral immune maturation of B cells into plasma cells through integrins, B-cell receptors, and TLRs (345). Leupaxin belongs to the paxillin gene family of adaptor proteins, is recruited to podosomes in a kindlin 3 manner, and is believed to be responsible for the stability and longevity of podosomes (346, 347) Podosomes are integrin/actin-rich blebs that appear on the surface of cells and play a role in dynamic cellular mobility, not only in the immune system but also in the literature as they are involved in the proliferation and migration of diverse cancer cells (348).

### ATF3 and 4

LPS tolerance involves severe loss of transcription of Aft3 and Atf4. Bioinformatics data mining has revealed losses of ATF3 in various human cancers such as HCC tumor tissues, corresponding to advanced clinical/pathological grade-stage (349) where elevated levels of ATF4 are linked to Treg cells embedded within tissues associated with tumor growth (350). There was also a tolerance nonresponse of *Gbp2 (−2.99 Log2FCp-VAL<1.00E-300) *Gbp2b (−3.25 Log2FC p-VAL 3.00E-306), and *Gbp5 (−2.75 Log2FC p-Val 2.37E-30) which are associated with poor overall survival of skin cancer patients (351). The data revealed chronic inflammatory losses in serglycin (Srgn). Srgn belongs to the family of proteoglycans with varying functions by cell type with known controlling elements over processes involved in the secretion of cytokines and granules by macrophages (TNF was also reduced in this study), (352) NK cells (perforin), mast cells (histamine, chymase, tryptase, carboxypeptidase), neutrophils (elastase) and (cytotoxic T lymphocytes CTLS granzyme B (storage)) (353, 354). SRGN-deficient (SG (−/-)) mice, display faulty antitumor/viral CD8+ T-cell and NK functions, reduced CD4(+) cells, reduced platelet function, and clotting, increased CD45 RC(+) leukocyte populations, enlarged lymphoid tissues, and poor prognoses in cancers (355–357) and viral infections (358) (354). Our work is in agreement with the observations of Zernichow *et al*. (2006), who reported parallel losses in SRGN to profile losses in the TNF response, during chronic inflammation-mediated tolerance.

Owing to tolerance, the expression of this transcript (SERPINB2) was attenuated during the chronic period but was still elevated above baseline compared with that at rest. SerpinB2 (plasminogen activator inhibitor 2) is a major product of activated monocytes and macrophages and is substantially induced during most inflammatory processes. Moreover, members of this family of serine protease inhibitors can block serine proteinase urokinase-type plasminogen activator (uPA) (359), which, if elevated, is involved in resistance to cisplatin-based chemotherapy (360), is indicative of aggressive oral squamous cell carcinoma (361) and is subject to polymorphisms linked to a range of inflammatory diseases involving adaptive immunity (362).

### Losses in Detoxification Systems

Chronic inflammation is associated with the loss of transcripts involved in glutathione conjugation/detoxification which could be involved in vulnerability to cancer initiation and/or growth in established cancers, such as glutathione S-transferase mu 1 (GSTM1) (363), SOD1,2, Srxn1 and Ephx1 (microsomal epoxide hydrolase is involved in the detoxification of environmental pollutants such as polycyclic aromatic hydrocarbons) (364). Importantly, our work also revealed that the elevation of the P450 (cytochrome) oxidoreductase (POR) gene is sustained during acute and chronic inflammation, creating paradoxical forces that require further study.

### S100s

There are 24 identified S100 proteins that participate in Ca^2+^ signal transduction of diverse natures and are involved in receptor for advanced glycation end products (RAGE), and only a few of these proteins are involved in TLR4 signaling. The data in this study showed that only S100A8 was minimally expressed in resting cells and was activated by LPS via the TLR4 receptor, with S100A1, A4, and A6 all being reduced (in the opposite direction) with acute and chronic inflammation. Although, S100 proteins are known to be expressed in macrophages, they are extremely diverse in nature in terms of function (365, 366). S100A1 is an inhibitor of protein kinase C signaling, the loss of which is associated with hypertrophic growth in heart tissue; S100A4 is involved in B-catenin cell motility and tumor invasion; and E-cadherin, B-catenin and S100A6 are involved in cell proliferation, where S100A8, is inducible by proinflammatory stimuli and defends against NO (181) (see Review (366). While there has been research on the consequences of S100 proteins, our work aligns in part with those reporting some type of connection between concomitant losses of S100a4 and CCL5 (RANTS) (367)/colony-stimulating factor. Homozygous S100A4(−/-) mice display impaired recruitment and chemotactic infiltration of macrophages to sites of inflammation in vivo (368). The loss of CCL5 in this study, upon chronic inflammation was notable because it was the only chemokine subject to tolerance, with full nonresponse *CCL5 (−8.83 Log2FCp-VAL 3.00E-303). Although the bulk of the literature on CCL5 suggests a role in driving tumor growth, our work supports the few studies reporting otherwise, showing higher CCL5 concentrations tied to a positive clinical outcome, along with IL1a (also suppressed in this work) (369), with amplification of the CCL5/CCR5 axis in colon cancer leading to the development of antitumor dendritic cells and CD8+ cytotoxic T cells (370, 371).

Chronic inflammation leads to the loss of the TYRO protein tyrosine kinase-binding protein, an adapter protein that, in humans is encoded by the TYROBP gene which is involved in the activation of multiple immune cells, including T cells, B cells, and macrophages. The low expression of TYROBP may promote the occurrence and progression of cancers and according to the risk model, lower TYROBP expression is associated with a greater risk of aggressive cancers (372) and lower survival rates in osteosarcoma patients (373).

Last, the category of reverse tolerance, is indicative of high levels during chronic inflammation that are no longer repressed (acute), the most significant being the elevation of PTMA prothymosin alpha, a biological response modifier (alarmin) known for its association with lipid droplets (LDs) inherent to colon cancer tissue involved in chemoresistance to gemcitabine, and elevated in esophageal cancer (EC) tumor tissue vs adjacent normal tissue (374, 375), a new target for therapeutic/diagnostic approaches in immune-related diseases such as cancer, inflammation, and sepsis (376).

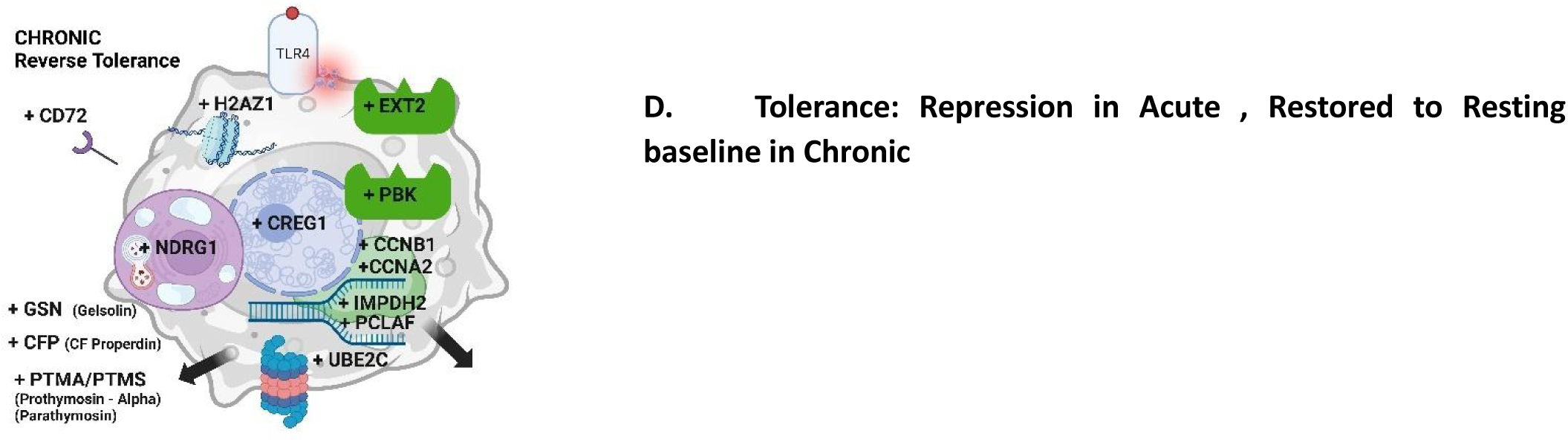

## Conclusion

The data in this study provide evidence to support the notion that tolerance and loss of immunosurveillance are associated with chronic inflammation. Moreover, we provide details on the phenomenon of tolerance as it could pertain to the limited efficacy in human cancers with repeated administration of synthetic lipoglycans or TLR4 agonists.

## Contributions

E.M. was involved with concept, experimental design, data analysis, bioinformatics, experiments,image design and manuscript preparation and literature review. A.B. assisted in carrying out all experiments needed including method development / design and validation. R.B. was involved with daily trouble shooting and steering of experimental design. S.D.R was involved with manuscript preparation/ review and K.FA S. saw to operational strategy and oversight of the entire project to manuscript submission.

## Acknowledgment

We thank Zhiyi Liu - LC Sciences for sequencing services and Steve Council of Gnabre Research Institute for their assistance with this review of the manuscript.

## Data Sharing

The data are available from the National Institutes of Health (NIH) Gene Expression Omnibus; series number GSE263901.

